# *eOmics*: an R package for improved omics data analysis

**DOI:** 10.1101/2023.03.11.532240

**Authors:** Yu Liu

## Abstract

Many computational tools have been developed for high-throughput omics data, and some are very popular, such as *limma, WGCNA*, and *EnrichR*. However, they also exhibit disadvantages in some special cases, such as imbalanced data analysis, causal inference, gene network functional enrichment, etc. Hence, we developed the R package *eOmics* to provide a comprehensive pipeline with these problems addressed. It combines an ensemble framework with *limma*, improving its performance on imbalanced data. Moreover, it couples a mediation model with *WGCNA*, so the causal relationship among *WGCNA* modules, module features, and phenotypes can be found, and this model is also used to explore the relationship between different omics. In addition, our package has some novel functional enrichment methods, capturing the influence of topological structure on gene set functions. Finally, it contains multi-omics clustering and classification functions to facilitate machine-learning tasks. Some basic functions, such as ANOVA analysis, are also available in it. The effectiveness of our package is proved by its performance on the three single or multi-omics datasets here. *eOmics* is available at: https://github.com/yuabrahamliu/eOmics.

## INTRODUCTION

High-throughput omics technologies provide unprecedented opportunities for mining biological information from data. Correspondingly, many methods have been developed for these tasks, and several have become very popular.

For example, *limma* has been used as a classic tool to identify differential features between sample groups because its Bayesian strategy borrows information between features, providing reliable inference even with small sample sizes (1). *WGCNA* is widely used to explore correlation patterns among genes, and its soft thresholding method preserves the continuous nature of the correlation, avoiding information loss and enhancing the robustness of the result (2). In addition, some gene functional enrichment tools, such as *DAVID* and *EnrichR*, are used downstream to check the functions of selected features (3,4). These tools constitute a general workflow for omics data analysis.

However, when using them practically, many factors can influence their performance. For example, when calling differential features, it is common to meet sample groups with imbalanced sizes, especially in clinical studies, which can impair the power of *limma*. Although *WGCNA* can find the gene modules correlated with a phenotype, this correlation is undirected, so it is unclear whether the module causes the phenotype or *vice versa*. In addition, when annotating the module functions, *DAVID* and *EnrichR* use the nodes of a module as inputs. However, this process does not consider the module’s edges, which reflect the module’s structure. Moreover, because different genes and gene-gene interactions have different importance, their weights should also be included in the analysis. Otherwise, any modules with the same nodes will have the same function annotation, even if their structures are different. Finally, as multi-omics data become more and more common, the relationship among different omics needs to be addressed, but the tools above mainly handle single-omic data without considering this relationship.

Hence, we developed the R package *eOmics* (enhanced omics) to address these computational challenges. Within the package, we combined an ensemble framework with *limma* to improve its performance on imbalanced data. In addition, we developed a mediation analysis model to explore the causal relationship between *WGCNA* modules and phenotypes, and this model was also used to find the relationship between different omics. We also considered gene-gene interactions and their weights for functional enrichment to account for the gene set structures. Furthermore, in virtue of some multi-omics techniques, such as ensemble and CCA (canonical correlation analysis), we made the package suitable for implementing clustering and classification on multi-omics data. Other basic functions, such as ANOVA analysis, missing value imputation, and dimension reduction, can also be found in the package.

To test the performance of *eOmics*, we applied it to three single or multi-omics datasets. Results show that it performs well on differential feature identification, causal inference, multi-omics classification, etc. We believe it can provide more possibilities for exploring omics data.

## METHODS AND RESULTS

### Package overview

The package contains five modules (Figure 1). The first is the ensemble-based *limma* module. It integrates *limma* and SMOTE (synthetic minority over-sampling technique) into a bagging framework, improving the power of *limma* for imbalanced sample groups.

**Figure 1.**
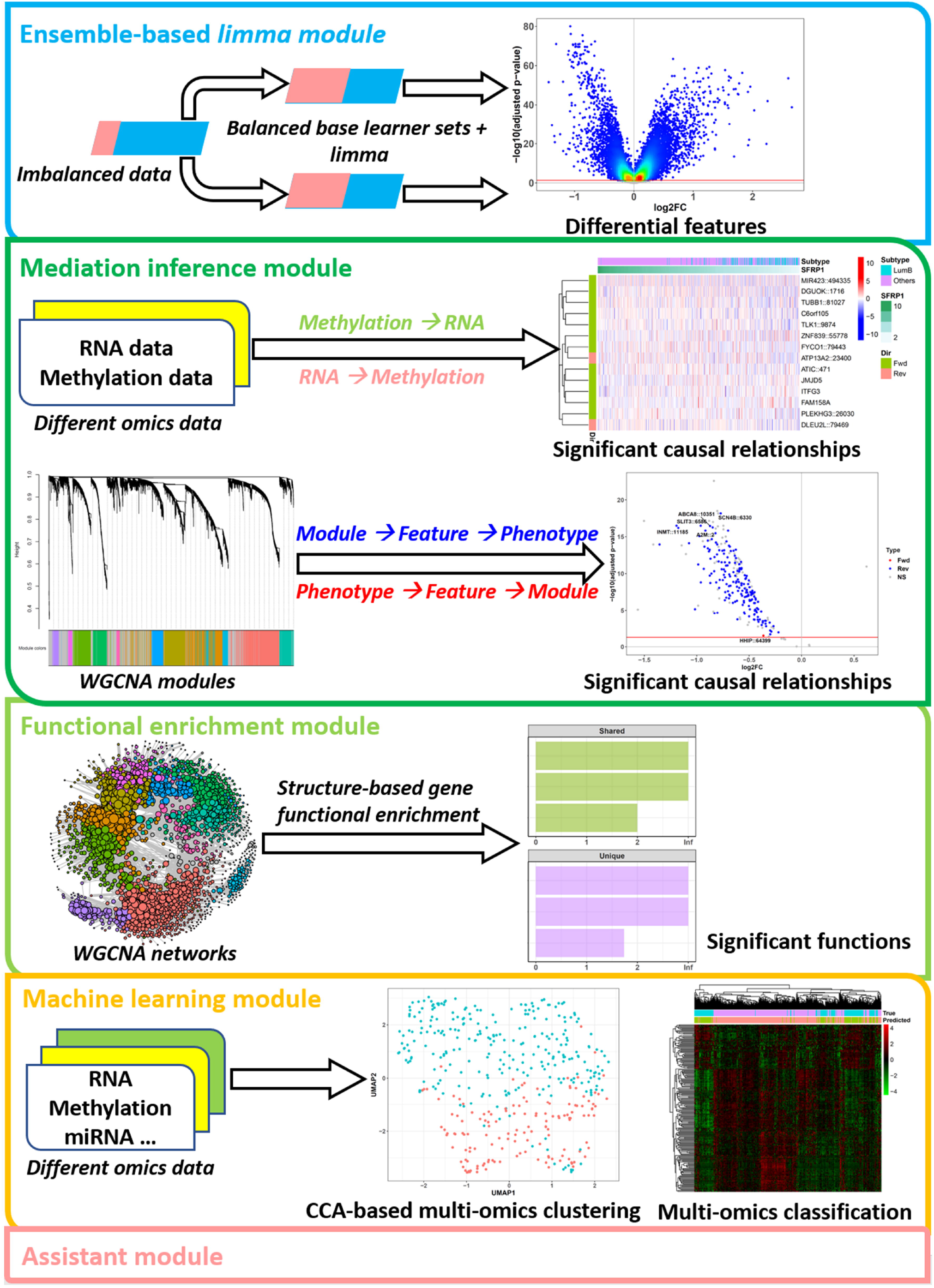
Package modules. The package has five modules: the ensemble-based limma module, the mediation inference module, the functional enrichment module, the machine learning module, and the assistant module.

The second is the mediation inference module. It inserts mediation analysis into the *WGCNA* pipeline, revealing the causal relationship among phenotypes, *WGCNA* modules, and module features. In addition, it is also combined with LASSO and multi-omics analysis, so the causal relationship between different omics features can be found.

The third is the functional enrichment module. It uses a correlation-based method to convert the analysis on non-RNA omics to RNA, reducing noise in the original data. Moreover, this module also performs structure-based enrichment, considering the gene pairs and their weights in any gene network, so the topological influence on gene set function can be captured.

The fourth is the machine learning module. To cluster samples from multi-omics, it applies a CCA-based method to compress the data. Then, *k*-means is used on this compressed one to cluster the samples. It also performs multi-omics classification, constructing a model with elastic net and an ensemble framework, which integrates different omics and aggregates their results into a final one. In addition, this module is closely related to *WGCNA* because it can also cluster with the *WGCNA* modules called from various omics.

The final is the assistant module. It provides various functions, such as missing value imputation, ANOVA analysis, and methylation site annotation.

The details of these algorithms can be found in Supplementary Data.

### The package performs well on placenta DNA methylation data

To test the performance of *eOmics*, we applied it to three single or multi-omics datasets. The single-omic one was a human placenta DNA methylation (DNAm) dataset collected from various studies. After data preprocessing and batch adjustment, we only kept the probes with high quality and were covered by all the Illumina 27K and 450K datasets. The final contained 18626 probes and 359 samples, including 258 normal samples and 101 disease samples with preeclampsia pregnancy complications caused by placental abnormalities.

The metadata of the 359 samples covered 4 phenotypic variables, including sample group (preeclampsia or control samples), gestational week, baby gender, and maternal ethnicity. Then, we checked their relationship to the methylation beta values using the function *featuresampling* in our package. For each phenotypic variable here, this function performed type-□ ANOVA to calculate its averaged F statistic across the top 10000 most variable DNAm probes, representing the variance it explained for the dataset. The result showed that the sample group, gestational week, and baby gender accounted for the most variance (Figure S1). It implied that: 1) this dataset was suitable for checking the methylation difference between the sample groups, and 2) but variance from the gestational week and baby gender needed to be removed. They were confounding factors for preeclampsia analysis, especially gestational week, which was strongly related to preeclampsia because this disease led to preterm delivery, so its samples always had small gestational week values (5). Hence, we used our function *difffeatures* to call the differential DNAm probes between preeclampsia and control groups. It could perform normal *limma* analysis with gestational week and baby gender adjusted.

However, the dataset had a much larger control group than the preeclampsia one (258 samples/101 samples), and this imbalance would impair *limma*’s power. To solve this, we switched *difffeatures* to another mode, which implemented ensemble-based *limma*. As described in Supplementary Data, it used up-down sampling to generate several base learner datasets with balanced groups, i.e., a large group would be down-sampled, and a small one would be up-sampled with the SMOTE method, so a final set with balanced groups would be generated. This process would be conducted several times to generate several such sets. Then, *limma* would be used on each of them, and their results would be ensembled to get the final (Figure S2).

In addition to up-down sampling, the balanced base learner sets could also be generated with up-sampling and down-sampling, respectively. Hence, we compared their performance using the simulated experiments described in Supplementary Data. The result showed that *limma* with up-down sampling called the differential DNAm feature set closest to the true one (Figure S3A). Although its precision and recall were not the best, after combining these metrics, its final F1 statistic was. Moreover, if controlling the false positive features more strictly, the F1 statistic became F0.5 by assigning a higher weight to precision, and it still showed that *limma* with up-down sampling performed better than normal *limma* and other methods.

Hence, we used the up-down sampling mode to implement ensemble-based *limma*. On the placenta dataset, it called 473 hyper and 1956 hypomethylated DNAm sites for the preeclampsia group (Figure 2A), more than that called by the normal *limma* method (Figure 2B). Further comparison showed that all the 1418 hypomethylated sites called by normal *limma* were included in the 1956 up-down sampling ones, and 272 of the 276 hypermethylated sites from normal *limma* were also called by up-down sampling *limma* (Figure 2C).

**Figure 2.**
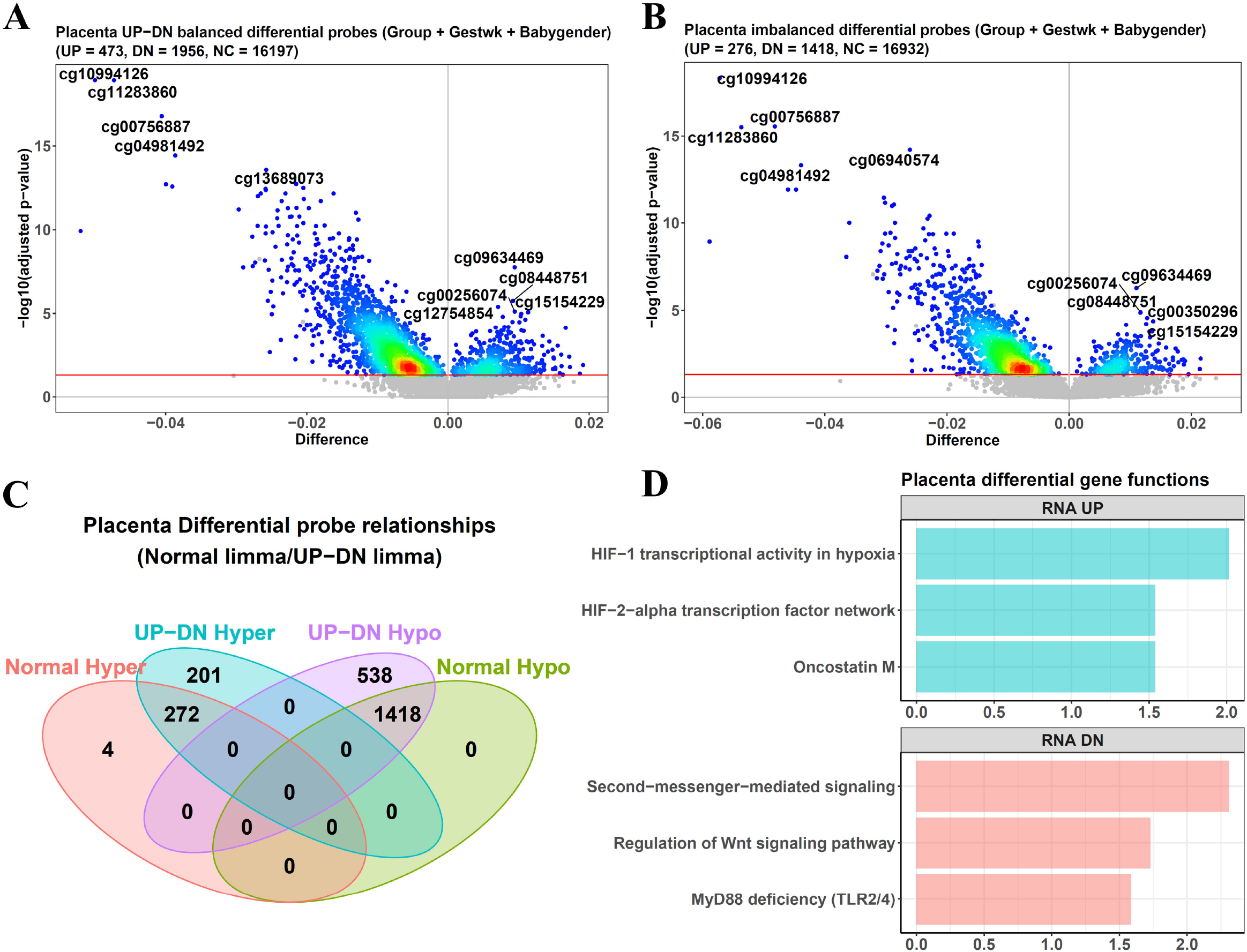
Ensemble-based limma performance on placenta DNA methylation data. (A) and (B) Ensemble-based limma (A) calls more differential DNAm probes from the placenta data than imbalanced (normal) limma (B). The probes with an adjusted p-value < 0.05 are called and represented as colorful dots, but a few dots located in confounding relevant DMRs (DNA methylation regions) will be filtered out and still gray. (C) Comparison between the differential DNAm probes called by up-down sampling limma and normal limma. (D) Correlation-based functional enrichment for the DNAm features. The x-axis represents −log10(adjusted p-value).

Besides, *difffeatures* could further filter these sites with its parameter *removereg*, which accepted genomic regions related to confounding factors and filtered out any sites there. All the analyses above included this step and used confounding DMRs (DNA methylation regions) as the genomic regions, which were called from several public 450K placenta datasets using the function *sigdmr* in our package. However, because the regression step of *limma* had adjusted the confoundings once, we only found 5 sites that needed this filtering, so its effect was limited and unnecessary, and this step could be skipped.

After calling the DNAm sites, another function, *corenrich*, was used to find their enriched functions. It utilized paired RNA-DNAm data to identify genes with an RNA expression significantly correlated to the differential DNAm sites and used them to perform functional analyses. Compared with the traditional methods directly on hyper and hypomethylated genes, this correlation step removed those whose DNAm change has little influence on their gene expression and avoided their misleading effects.

Our placenta dataset contained 48 samples (30 preeclampsia and 18 control samples) with paired RNA microarray data, which were used by *corenrich* to find the correlated genes to apply this method, and the result was well related to the disease (Figure 2D). For example, it identified the function “HIF-1 transcriptional activity in hypoxia” as prompted in preeclampsia. On the other hand, this disorder was always linked to an elevated level of HIF-1α, a critical regulator of placenta development (6). Also, “Oncostatin M” was revealed as enhanced in the disease, and “Regulation of Wnt signaling pathway” was found as down-regulated, consistent with previous experimental conclusions (7,8).

Next, we performed *WGCNA* analysis on the data. We first used our function *probestogenes* to compress the DNAm probe beta values to genes, and then another function, *diffwgcna*, was used. At first, we also tried combining *WGCNA* analysis with the up-down sampling ensemble framework and aggregating the base learner results via the consensus *WGCNA* algorithm (9,10). However, our simulated experiments showed that this method could not improve *WGCNA*’s performance on the F1 and F0.5 values (Figure S3B). Hence, *diffwgcna* only used the normal *WGCNA* method to call modules, and on the placenta dataset, it found 11 (Figure S4A).

Then, we searched for the modules related to preeclampsia. Compared with the traditional *WGCNA* pipeline, which relied on correlation coefficient, *diffwgcna* used *limma* to find them. Because of the regression step of *limma*, it had the advantage of adjusting the confounding effects of the gestational week and baby gender, and at this *limma* step, the ensemble mode could be chosen by *diffwgcna*. The result showed that 7 of the 11 modules had a significant preeclampsia/control difference in their eigengenes (Figure 3A). Moreover, *limma* could also be used within the modules. For example, in the ME3 module, which showed the largest difference between preeclampsia and control groups, 417 of its 527 genes were found as hypomethylated in preeclampsia samples (Figure 3B).

**Figure 3.**
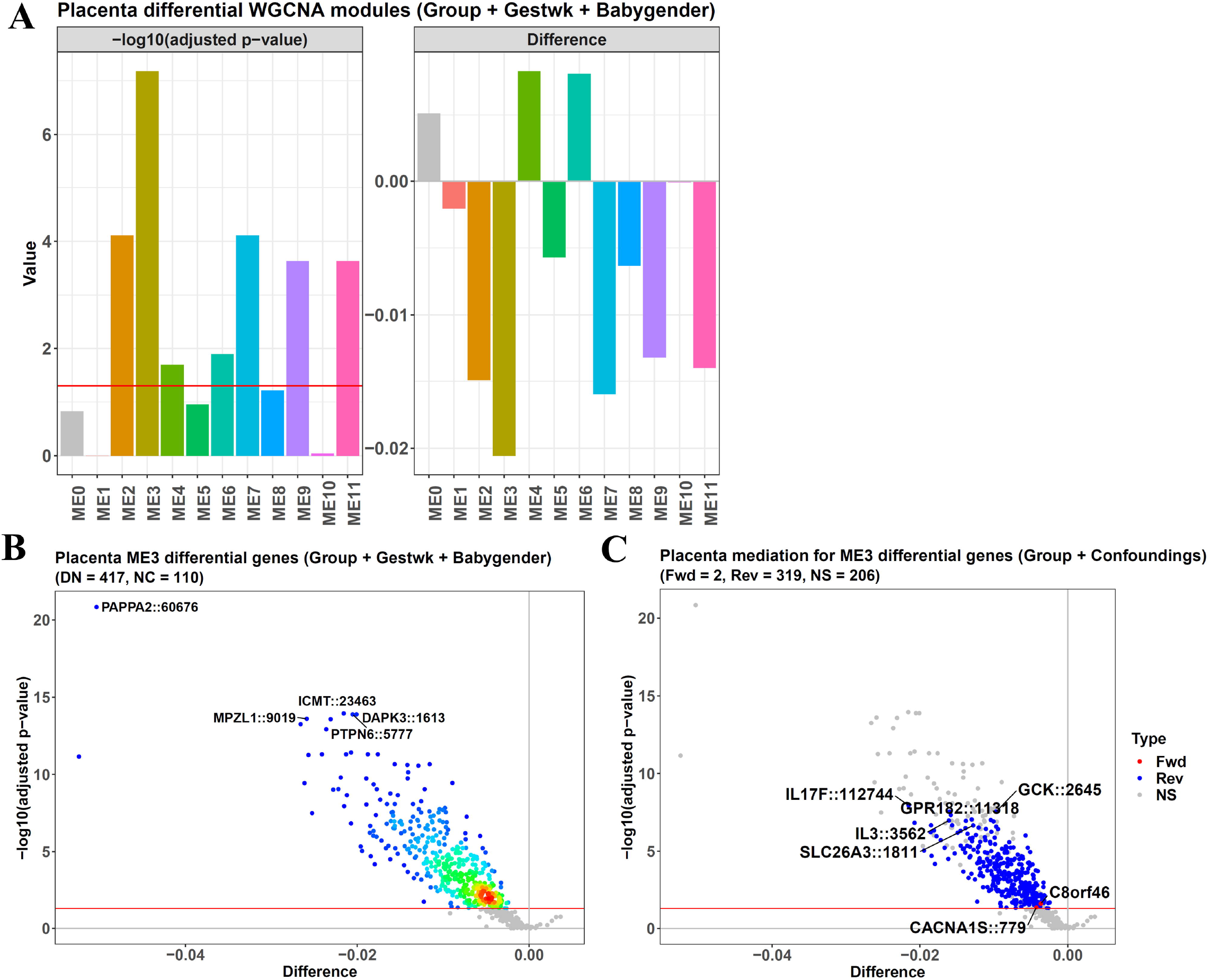
Mediation-coupled WGCNA performance on placenta DNA methylation data. (A) The diffwgcna function identifies 11 WGCNA modules, and 7 have largely different eigengenes between preeclampsia/control groups, as revealed by the limma method. (B) After conducting limma within the ME3 module, 417 of its 527 genes show significantly different beta values between the preeclampsia/control groups. (C) In the ME3 module, 2 of its genes mediate the relationship of “ME3→ME3 gene→preeclampsia” (the forward direction), and 319 mediate “preeclampsia→ME3 gene→ME3” (the reverse direction). The y-axis and x-axis show the −log10(adjusted p-value) and preeclampsia/control beta value difference when screening the differential genes with limma.

In addition, *diffwgcna* had another advantage of demonstrating the causal relationship among the phenotype, module, and module features. As described in Supplementary Data, it constructed mediation models for each differential feature in a module, using phenotype status, module eigengenes, and feature values. Then, the models could test 2 causal directions, and in the ME3 module here, this analysis found that 2 of its genes mediated the direction of “ME3→ME3 gene→preeclampsia”, and 319 genes mediated the opposite of “preeclampsia→ME3 gene→ME3” (Figure 3C). Hence, the disease caused the module changes mostly, and the module drove the disease rarely.

The 319 genes showed an association with inflammatory stress in preeclampsia, such as IL3, IL17F, and GPR182, which were the top genes mediating the effects of preeclampsia on ME3. Among them, IL3 and IL17F were cytokines for inflammatory responses (11,12), and GPR182 was a G protein-coupled receptor interacting with chemokines CXCL10, 12, and 13 (13). Because these genes’ causal direction was from preeclampsia to ME3, the conclusion was that inflammation was not the cause, but the result, of this disease.

Meanwhile, CACNA1S and C8orf46 (VXN) were the only 2 genes mediating the causal direction from ME3 to preeclampsia. CACNA1S had expression in the placenta (14,15), and encoded the α-1S subunit of the dihydropyridine receptor, a calcium channel responsible for vascular contraction. Given that preeclampsia was a pregnancy hypertension disease, the causal effect of CACNA1S on it was clear. Moreover, this gene was the target of amlodipine besylate, a small-molecule drug for preeclampsia treatment, further supporting that the inference here was reasonable. At the same time, the other gene, C8orf46, was largely unknown. However, it was still reported to work together with CDKN1B (P27KIP1) (16), a critical gene for mediating trophoblast invasion (17). On the other hand, it was generally agreed that insufficient invasion of placental trophoblasts was a driver of preeclampsia (18). Hence, C8orf46 might cause this disease by influencing trophoblast invasion.

Finally, we explored preeclampsia subtypes with our package. Its function *multiCCA* used CCA to convert multi-omics data into a compressed one and then performed *k*-means to get the clustering result. On the single-omic here, the CCA became PCA for the DNAm probe data of the 101 preeclampsia samples. We used the consensus clustering mode and tried different cluster numbers for the *k*-means step. From the 3 internal validation indices calculated by *multiCCA*, the best cluster number was 2 because it gave the maximum Silhouette and Calinski indices and the minimum Davies-Bouldin index (Figure S5A).

The 2 clusters had imbalanced sample sizes; the larger one (subtype1) contained 72 samples, and the smaller one (subtype2) contained 29 (Figure 4A). Hence, ensemble-based *limma* was used to find their differential DNAm probes, and it identified 1197 hyper and 13625 hypomethylated sites in subtype2 (Figure 4B). Then, *corenrich* checked the function of these sites. It used the former 48 paired RNA-DNAm samples to find the correlated RNA genes, then generated the enrichment result (Figure 4C). In subtype2, some immune functions were down-regulated, such as “Interleukin-1 regulation of extracellular matrix”. Hence, they were relatively up-regulated in subtype1, indicating its closer relationship with the inflammatory stress of preeclampsia. On the other hand, “Activation of TRKA receptors” was up-regulated in subtype2, and because TRKA stimulated HIF-1α expression (19), it had more connections with the oxidative stress of this disease.

**Figure 4.**
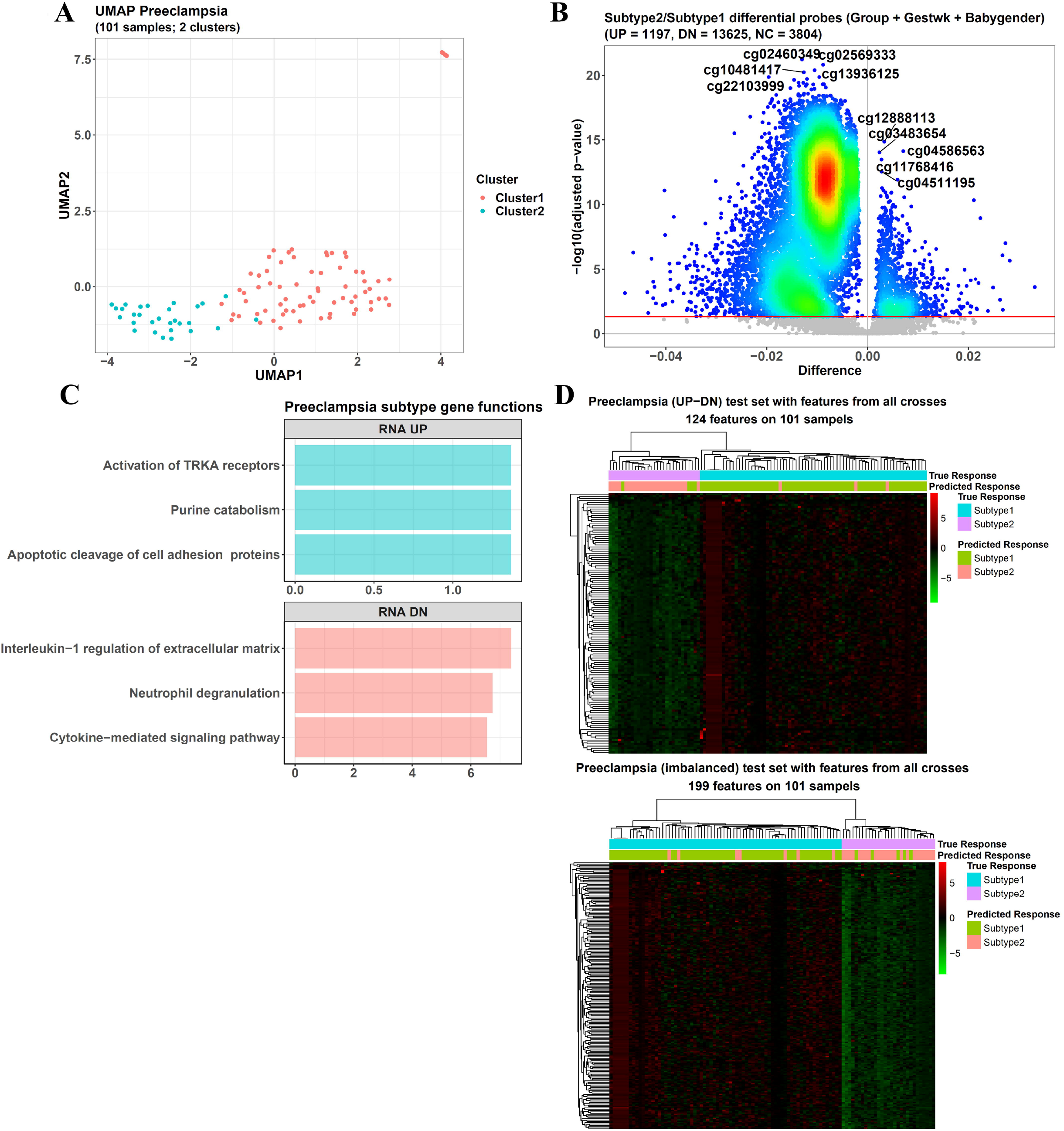
Package performance on preeclampsia subtype methylation data. (A) UMAP embedding of the 2 preeclampsia subtypes. (B) Differential DNAm sites between the 2 subtypes. (C) Correlation-based functional enrichment for the subtype2/subtype1 differential DNAm features. The x-axis is −log10(adjusted p-value). (D) The ensemble-based elastic net classifier (with up-down sampling) has an accuracy of 0.931 for the testing samples from all the 5-fold cross-validation sets, and that of the normal (imbalanced) elastic net model is 0.881. In the heatmaps, each column represents one sample, and each row represents one DNAm probe selected by the model. The entries are beta values after scaling across samples, and the hierarchical bars above the heatmap show the true sample labels and the predicted ones.

After getting the subtype labels, we trained a model to classify them with our function *omicsclassifier*. As demonstrated in Supplementary Data, it combined elastic net with the balanced ensemble framework to improve the performance on imbalanced data (Figure S6), similar to a previous method predicting drug responses (20). It could train a model from both single-omic data and multi-omics data. Here, the 5-fold cross-validation showed that it had a much better performance on the testing sets (accuracy = 0.931) than the normal elastic net model (accuracy = 0.881), which was also constructed by *omicsclassifier* (Figure 4D).

This case study demonstrated the advantages of *eOmics* in single-omic data analysis.

### The package performs well on lung cancer multi-omics data

Next, we applied the package on the TCGA LUAD (lung adenocarcinoma) dataset, which contained 419 cancer samples and 3 different omics: RNA, DNAm (450K), and miRNA. After preprocessing, their feature numbers were 21465 RNA genes, 412481 DNAm probes, and 1574 miRNA genes, respectively.

Then, we selected the top 12000 most variable genes and probes from the RNA and DNAm datasets. Together with the 1574 miRNA genes, they were transferred to *multiCCA* to perform multi-omics clustering. The 3 internal indices showed that the optimal cluster number was 2, with the largest Silhouette and Calinski indices (Figure S5B). Correspondingly, the first cluster (subtype1) contained 146 samples, and the second (subtype2) contained 273 samples. Most importantly, survival analysis showed a significant difference between them (log-rank p-value = 0.0353), with subtype1 having a longer survival time (Figure 5A). In contrast, using any single-omic for *multiCCA* clustering could not generate subtypes with significant survival difference (Figure S7A to C), indicating the advantage of multi-omics. In addition, the package also contained another function, *wgcnacluster*, which performed *WGCNA* on each omic first and then combined their *WGCNA* eigengenes for multi-omics clustering. However, its subtypes also had no significant survival difference (Figure S7D). Hence, the multi-omics subtypes from *multiCCA* were chosen for the following analysis.

**Figure 5.**
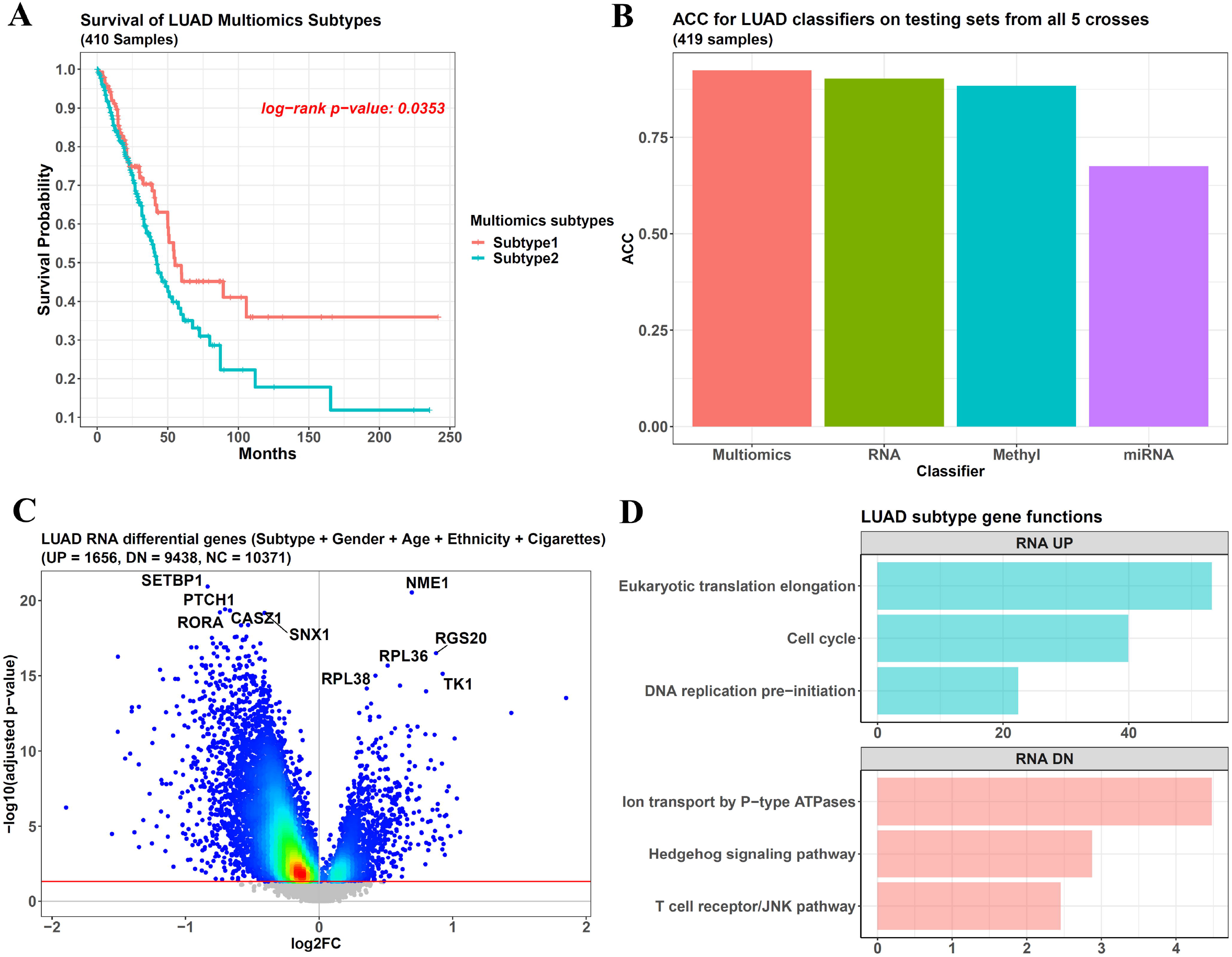
Package performance on LUAD multi-omics data. (A) The 2 LUAD subtypes from multiCCA multi-omics clustering show a significant survival difference. Because only 410 samples of the original 419 ones have survival information in TCGA, only they are included here. (B) The testing set accuracy of the omicsclassifier models from multi-omics, RNA, DNAm, and miRNA data after 5-fold cross-validation (0.924, 0.902, 0.883, and 0.675, respectively). (C) Ensemble-based limma calls differential genes from the LUAD RNA data and finds 1656 up-regulated and 9438 down-regulated genes in subtype2 compared with subtype1. The genes with an adjusted p-value < 0.05 are called and represented as colorful dots. (D) Functional enrichment for the up and down-regulated genes in LUAD subtype2. The x-axis represents −log10(p-value).

Then, we used *omicsclassifier* to classify these subtypes and found that for a 5-fold cross-validation, utilizing all the 3 omics could reach an accuracy of 0.924 on the testing set, higher than using any single-omic (0.902, 0.883, and 0.675 for single-RNA, DNAm, and miRNA data) (Figure 5B), showing the advantage of multi-omics classification.

Next, we compared the subtype gene expression by applying *difffeatures* to their RNA data, and given their imbalanced sample sizes (273 subtype2/146 subtype1), we used ensemble-based *limma*. According to the ANOVA analysis from *featuresampling*, the confounding factors here were patients’ gender, age, ethnicity, and cigarettes used per day, and all of them had an F statistic > 1 (Figure S1B). Finally, 1656 genes were detected as up-regulated in subtype2, and 9438 were down-regulated (Figure 5C). Then, we used *corenrich* to check their functions, and because the features had already been RNA genes, its correlation step for omics conversion was skipped. The results showed that the up-regulated genes in subtype2 were enriched in “Cell cycle” and “DNA replication pre-initiation”, etc., and some tumor-suppression functions, such as “T cell receptor/JNK pathway”, were down-regulated, explaining the weaker survival of subtype2 (Figure 5D).

Then, we used *diffwgcna* on the RNA data to identify *WGCNA* modules related to the subtype difference. It detected 11 modules (Figure S4B), and 10 were significantly related to the subtypes (Figure 6A). The ME8 module showed the closest relevance, and *limma* within it found 250 RNA genes as largely down-regulated in subtype2 compared with subtype1, with only 1 gene up-regulated (Figure 6B). Further mediation analysis identified 1 gene mediating the causal direction of “ME8→ME8 gene→LUAD subtypes” and 176 genes mediating the opposite direction of “LUAD subtypes→ME8 gene→ME8” (Figure 6C).

**Figure 6.**
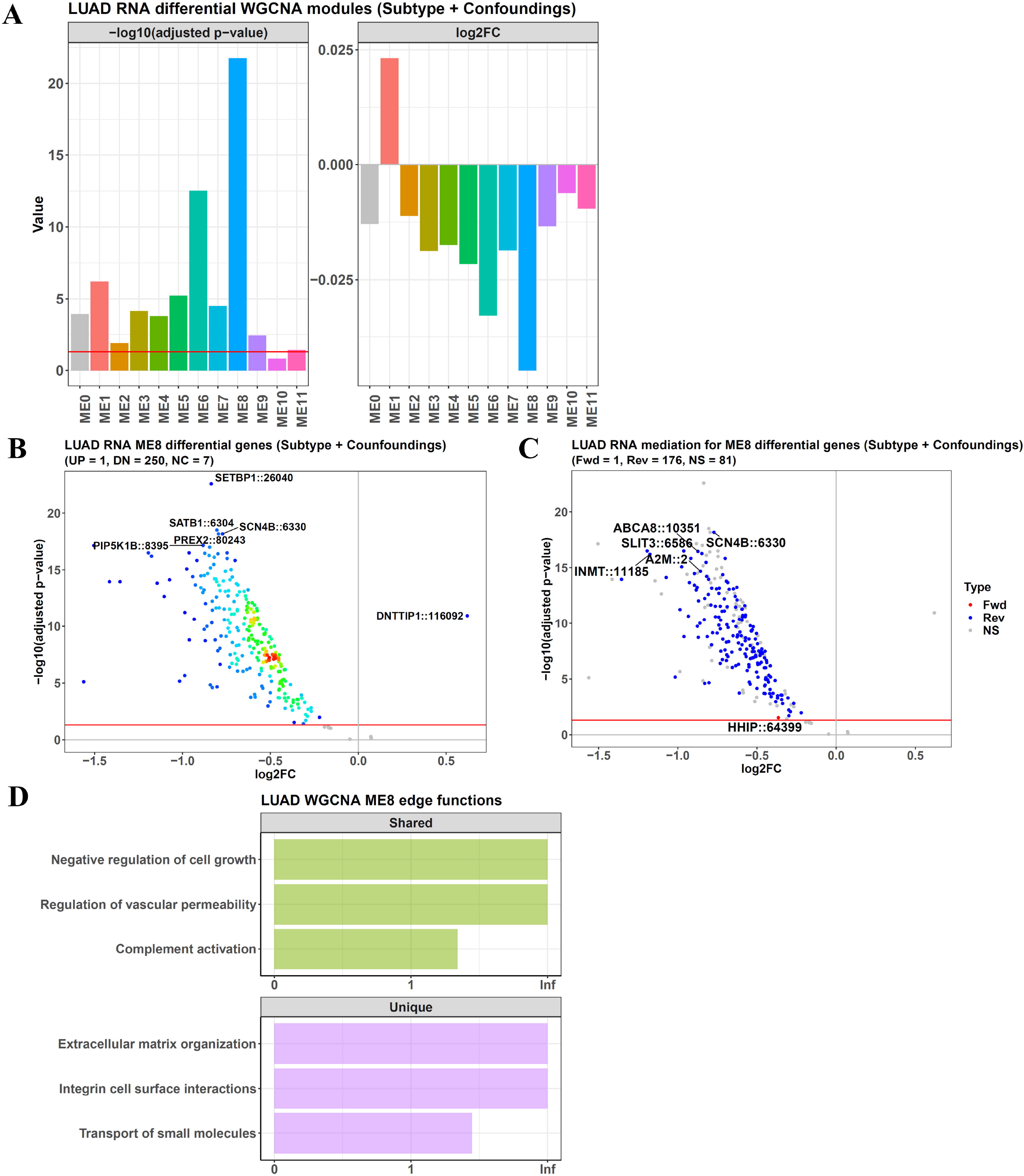
Performance of diffwgcna on LUAD RNA data. (A) The function diffwgcna detects 11 modules in the LUAD RNA dataset, and 10 have significantly different eigengenes between subtype2/subtype1 groups, as revealed by ensemble-based limma. (B) In the ME8 module, ensemble-based limma finds that 251 genes have significantly different expressions between the 2 LUAD subtypes. (C) Mediation analysis finds 1 gene in ME8 mediates the causal direction of “ME8→ME8 gene→LUAD subtypes” (the forward direction), and 176 mediate “LUAD subtypes→ME8 gene→ME8” (the reverse direction). The y-axis and x-axis show the −log10(adjusted p-value) and subtype2/subtype1 RNA expression difference when screening the differential genes with limma. (D) Edge-based functional enrichment for the genes in ME8. The green bars are functions identified by both the edge-based method and the traditional gene count method, and the purple bars are only found as significant by the edge method. The x-axis represents −log10(adjusted p-value) in the edge-based method.

The 1 gene for the first direction was HHIP, a conserved hedgehog-interacting protein able to modulate this signaling. Because the hedgehog pathway was highly associated with cancer stemness (21,22), the causal relationship here indicated that cancer stemness was the cause for LUAD subtype divergence.

In contrast, for the 176 genes in the opposite direction, their changes were the results of this subtype divergence. The top genes, such as SCN4B and SLIT3, controlled cancer cell migration (23,24), and the ABCA8 and A2M genes were related to cancer chemoresistance and hemostasis, respectively (25,26).

Furthermore, *diffwgcna* checked the overall function of ME8 with structure-based functional enrichment. It was different from traditional gene enrichment by considering the modules’ network structure, reflected by the gene-gene interaction edges and edge weights, and this was more reasonable because a gene needed to interact with others to function. As illustrated in Supplementary Data, the function terms of each edge (gene-gene pair) were defined as the intersection of its 2 genes’. Then, an edge weight shuffling-based method was used to find the function terms with significantly large weights.

For ME8, this method found that it covered some cancer-relevant functions such as “Negative regulation of cell growth” and “Regulation of vascular permeability” (Figure 6D). Meanwhile, they were also identified by the traditional gene count enrichment, which could be conducted by another function, *topoenrich*, in the package. In contrast, some functions were only significant in the edge-based method, such as “Extracellular matrix organization” and “Integrin cell surface interactions”. They were insignificant in the traditional method because their gene counts were under the significant cutoff of the enrichment test. However, their edge weights were unusually large so that they could be identified by the edge method.

Although this structure-based enrichment was achieved by shuffling, it could also be fulfilled via a Γ function-based hypergeometric test, but because applying it to the fully-connected *WGCNA* network brought a huge computational burden, *diffwgcna* only covered the top 1% weighted edges. These methods could also be called with the function *topoenrich* in our package. More details could be found in Supplementary Data.

This case study showed the application of *eOmics* in multi-omics data analysis.

### The package performs well on breast cancer multi-omics data

We also tested the package using 752 cancer samples in the TCGA BRCA (breast invasive carcinoma) dataset, covering RNA, 450K DNAm, and miRNA omics. This time, we converted the DNAm probe values to gene beta values with our function *probestogenes*. After preprocessing, the feature numbers were 21502 RNA genes, 18648 DNAm genes, and 1582 miRNA genes. In addition, we collected the BRCA subtype information from *TCGAbiolinks* (27), which labeled the samples as 5 PAM50 subtypes (411 Luminal A samples, 137 Luminal B samples, 126 basal-like samples, 44 HER2-enriched samples, and 34 normal-like samples).

Then, we used *omicsclassifier* to classify the sample subtypes. For the RNA and DNAm data, their top 10000 most variable genes were used, and for the miRNA data, all the 1582 miRNA genes were used. For the classifier using the 3 omics together, its 5-fold cross-validation accuracy on testing data was 0.890, higher than the single-omic ones (0.887, 0.834, and 0.814 for single-RNA, DNAm, and miRNA models) (Figure 7A). Hence, the advantage of multi-omics classification was shown again.

**Figure 7.**
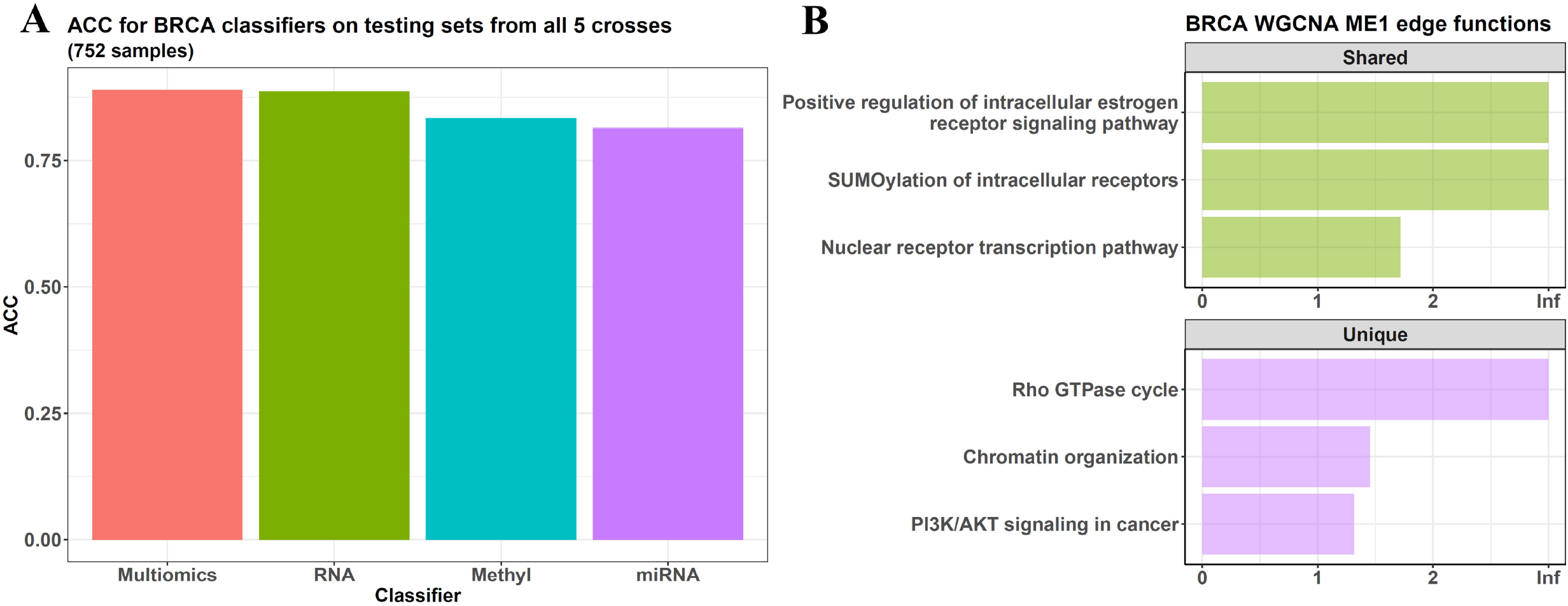
Package performance on BRCA multi-omics data. (A) When using RNA, DNAm, and miRNA data together, the function omicsclassifier can classify the BRCA subtypes during 5-fold cross-validation and generates an accuracy on testing data as 0.890, better than using only one omic (0.887, 0.834, and 0.814 for single-RNA, DNAm, and miRNA data, respectively). (B) Edge-based functional enrichment for the largest WGCNA module of the BRCA RNA data. The green bars are functions identified by both the edge-based method and the traditional gene count method, and the purple bars are only found as significant by the edge method. The x-axis represents −log10(adjusted p-value) in the edge-based method.

Next, *diffwgcna* was used on the RNA data to perform *WGCNA*. It found 13 modules (Figure S4C), and we tested the edge-based functional enrichment on the largest module, ME1 (containing 888 genes). The result showed that it was highly related to nuclear receptor functions, such as the “Nuclear receptor transcription pathway” and “Positive regulation of intracellular estrogen receptor signaling pathway”, which were also identified by the traditional gene count enrichment (Figure 7B). However, the edge-based method also found some pathways with significantly large edge weights, such as “PI3K/AKT signaling in cancer” and “Rho GTPase cycle”, which were missed by the gene count enrichment.

Then, we performed ANOVA on the RNA data to check the variances of the phenotypic variables and found that this dataset was largely related to the BRCA subtypes. However, the variance from patient ethnicity and age also had an F statistic > 1 (Figure S1C), and for the DNAm gene dataset, its ANOVA result was similar (Figure S1D).

Hence, we adjusted patient ethnicity and age when using *limma* to identify subtype-relevant RNA and DNAm genes. When comparing the Luminal A subtype with all other non-Luminal A samples, we found that it had 7426 genes over-expressed and 5928 under-expressed in the RNA data and had 6688 genes hypermethylated and 3209 hypomethylated in the DNAm data (Figure 8A and B).

**Figure 8.**
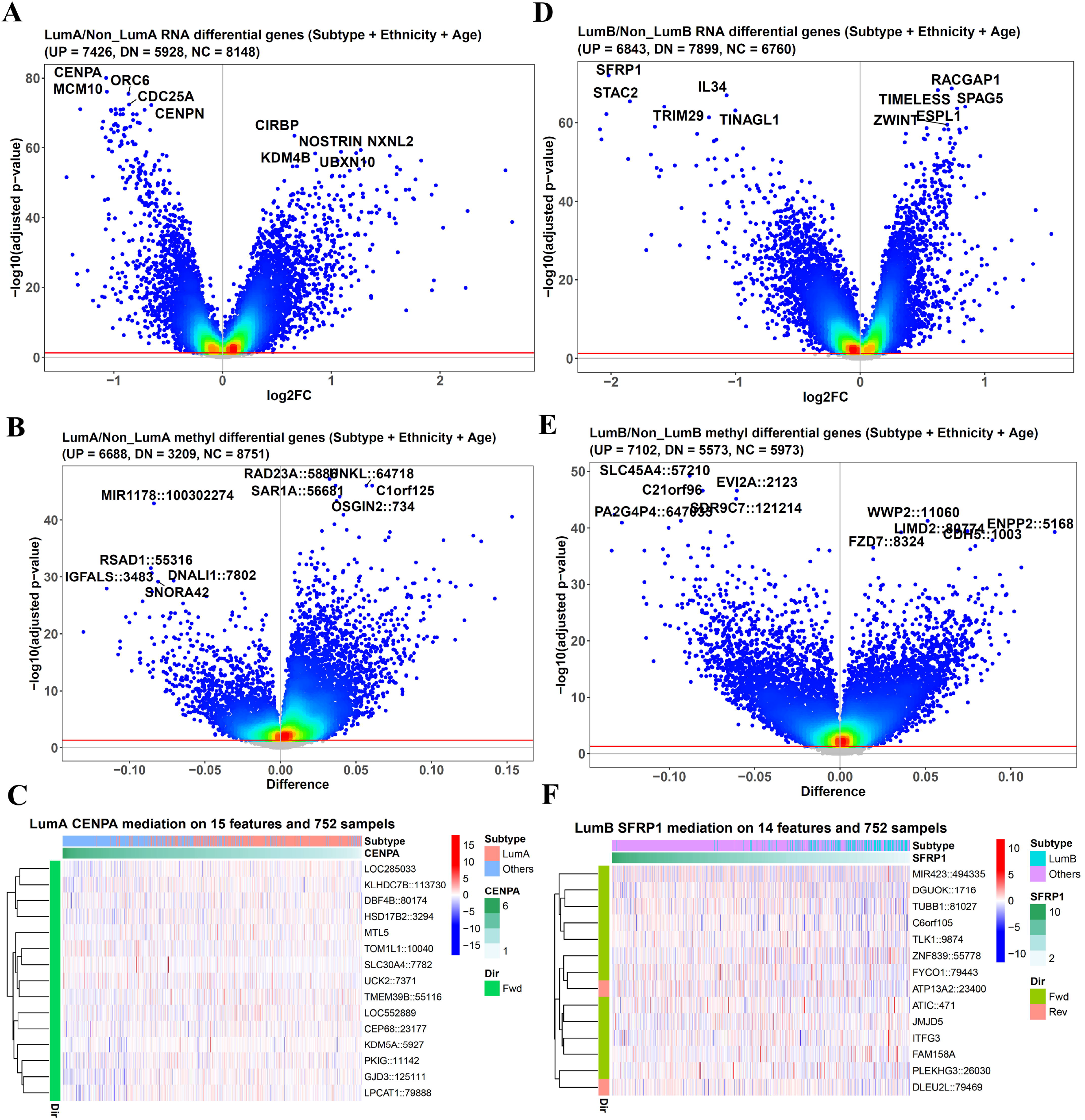
Multi-omics regulatory analysis on BRCA multi-omics data. (A) and (B) Ensemble-based limma finds differential RNA and DNAm genes between Luminal A subtype and non-Luminal A samples. (C) From the differential DNAm genes, lassomediation identifies 15 participating in the causal direction of “DNAm gene→CENPA expression→Luminal A/non-Luminal A” (the forward direction). The rows in the heatmap are these DNAm genes, and the columns represent samples ranked following their CENPA RNA values from high to low, as shown by the green hierarchical bar above the heatmap. The entries are beta values of the DNAm genes after scaling across samples. (D) and (E) Ensemble-based limma identifies differential RNA and DNAm genes in the Luminal B/non-Luminal B comparison. (F) For the RNA gene SFRP1, lassomediation finds 12 DNAm genes in the causal direction of “DNAm gene→ SFRP1 expression→Luminal B/non-Luminal B” (the forward direction) and 2 in the opposite of “Lumnal B/non-Luminal B→SFRP1 expression→DNAm gene” (the reverse direction).

Then, we explored the regulatory relationship between these RNA and DNAm genes, achieved by the function *lassomediation* in the package. As explained by Supplementary Data, it coupled the ensemble-based LASSO and the mediation models and tested the causal relationship among RNA gene, DNAm gene, and phenotype. For example, CENPA was the most differential RNA gene in the Luminal A/non-Luminal A comparison (adjusted p-value = 9.086e-81 and log2FC = −1.072), and from the differential DNAm genes above, *lassomediation* identified 15 participating in the causal direction of “DNAm gene→CENPA expression→Luminal A/non-Luminal A” (Figure 8C). Several of them had publication support, such as DBF4B, CEP68, and KDM5A.

CENPA itself was a histone H3-like protein in centromeric nucleosomes and propagated centromere identity through cell proliferation (28). On the other hand, *lassomediation* found DBF4B and CEP68 at its upstream. The former encoded the regulatory subunit for CDC7 and was the center of cell proliferation, and the latter was in charge of centrosome maintenance (29). Hence, their influence on CENPA’s expression became clear from their control roles in cell proliferation and centrosome. Furthermore, KDM5A was also identified as an upstream regulator of CENPA, consistent with the experimental conclusion that KDM5A could demethylate trimethylated Lys-4 of histone H3, and the result H3K4me2 could maintain CENPA (30,31). Thus, the causal relationship here clarified that in addition to regulating CENPA via H3K4me2, the methylation status of KDM5A could influence CENPA’s RNA expression.

We also checked the differential RNA and DNAm genes between the Luminal B and non-Luminal B samples and found 6843 up-regulated RNA genes and 7899 down-regulated ones. Meanwhile, 7102 genes were hypermethylated, and 5573 were hypomethylated (Figure 8D and E).

For the most differential RNA gene SFRP1 (adjusted p-value = 7.453e-73 and log2FC = −2.018), we performed the regulatory analysis and found 12 differential DNAm genes in the causal direction of “DNAm gene→SFRP1 expression→Luminal B/non-Luminal B” and 2 in the opposite direction (Figure 8F).

SFRP1 was an important modulator of Wnt signaling through direct binding with Wnts. This time, *lassomediation* identified TLK1 at its upstream, which encoded a serine/threonine kinase involved in chromatin assembly. Although no experimental study checked this TLK1-SFRP1 relationship directly, it was reported that the knockdown of TLK1 largely changed the Wnt pathway (32), and because of the direct binding between Wnts and SFRP1, it was reasonable to infer that TLK1 regulated SFRP1’s expression and then influenced Wnt signaling. In addition, 2 microtubule-relevant genes, TUBB1 and FYCO1, were also found upstream of SFRP1. Given that microtubule organization mediated Wnt activation (33,34), the causal relationship here could be understood, and it indicated that the cytoskeleton might also regulate Wnts by influencing their binding partner SFRP1.

This case study showed the performance of *eOmics* on multi-omics regulatory analysis.

## DISCUSSION

Many computational tools have been developed for analyzing high-throughput omics data. However, they always have some disadvantages when handling complex tasks in the real world. Hence, we developed the R package *eOmics* to overcome problems such as data imbalance, causal inference, multi-omics integration, etc.

We introduced an ensemble framework to *limma*, improving its power on imbalanced data. Meanwhile, our *WGCNA* pipeline contains a mediation analysis step, so the causal relationship among *WGCNA* modules, module features, and phenotypes can be revealed. This mediation model is also coupled with LASSO feature selection to explore the relationship between different omics, which is the central task of multi-omics analysis. In addition, our package includes novel functional enrichment methods, such as the edge-based one, considering the influence of topological structure on gene set functions. Finally, we added multi-omics clustering and classification functions to the package to facilitate machine-learning tasks.

These merits have been shown by the case studies above, but it is noteworthy that the traditional methods also have many advantages, so we included them in the package. For example, our simulated experiments showed that the ensemble-based *limma* called a differential feature set closer to the true one, with larger F1 and F0.5 statistics. However, if checking the precision only, the normal *limma* method showed an advantage over the ensemble, although its recall value was very weak, indicating many missed features during its imbalanced data analysis. Hence, if the purpose is to mine comprehensive information and a weaker precision is acceptable, ensemble-based *limma* will be suggested, but if it needs strict control over false positives, the normal one should be more suitable.

On the other hand, our edge-based enrichment can capture the topological influence on gene set functions, but its gene-gene combination can lead to a huge number of gene pairs, exerting much burden on the genomic background computation in hypergeometric or Fisher’s test. Thus, we recommend solving it with the edge weight shuffling-based method instead of these tests, largely reducing the computational costs. In this case, the edge background is the network where shuffling is performed, different from that of the hypergeometric test, which is the huge number of gene pairs in the genome. However, this also makes their results have different meanings: the hypergeometric one indicates what functions the gene set should have, and the shuffling one shows that among the gene set’s functions, which have the top priorities. Hence, the shuffling method cannot completely replace the enrichment tests.

Another important thing is that we emphasized confounding adjustment in all the case studies, and the function *featuresampling* was frequently used to perform ANOVA and determine the datasets’ confounding factors. It is especially important for differential feature analysis and mediation inference. The former needs to include the confoundings in its regression model to adjust their variance, and the mediation inference assumes that no unmeasured confounding exists for the relationship, and violations can give rise to misleadings (35–37).

Hence, confounding factors can influence the results’ correctness, and we attached great importance to it, with our package including several functions, such as *featuresampling* and *imputemeta*, to facilitate confounding analysis. However, in practice, there is always a possibility of unmeasured confounding. In this situation, biological knowledge becomes vital in judging the computational results. For example, in the LUAD subtype study, although the mediation model covered several confoundings, whether there were still missed ones was unknown, making its results not completely believable. However, from the genes’ biological functions, the HHIP gene was closely related to cancer stem cells, which were key drivers of tumor progression (38), so the inference that HHIP caused the subtype divergence became reasonable. It can be seen that biological knowledge coordinates the strict requirements on confounding factors and the difficulty of covering everything in practice. Hence, confounding information and biological knowledge should get enough attention when implementing differential and mediation analyses with our package. Their results need further curation via biological knowledge, publication searching, and sometimes experimental validation. In our package, we included some functions, such as *geneanno* and *probeanno*, to quickly provide some features’ biological information and facilitate this curation, but for the real experimental validation, it emphasizes and relies on the work of experimental researchers.

In summary, we developed the R package *eOmics*. It includes novel functions to handle tasks on imbalanced data, causal inference, functional enrichment, etc. We hope it can help researchers explore the biological information underlying high-throughput omics data.

## Supporting information

Tutorial

## SUPPLEMENTARY DATA

*eOmics* is available on GitHub (https://github.com/yuabrahamliu/eOmics). Its tutorial can be found in the supplementary files or at https://github.com/yuabrahamliu/eOmics/blob/main/ README.md. This paper also has other Supplementary Data available.

## REFERENCES

1. Ritchie, M.E., Phipson, B., Wu, D., Hu, Y., Law, C.W., Shi, W. and Smyth, G.K. (2015) limma powers differential expression analyses for RNA-sequencing and microarray studies. Nucleic Acids Res, 43, e47.

2. Zhang, B. and Horvath, S. (2005) A General Framework for Weighted Gene Co-Expression Network Analysis. Statistical Applications in Genetics and Molecular Biology, 4.

3. Sherman, B.T., Hao, M., Qiu, J., Jiao, X., Baseler, M.W., Lane, H.C., Imamichi, T. and Chang, W. (2022) DAVID: a web server for functional enrichment analysis and functional annotation of gene lists (2021 update). Nucleic Acids Research, 50, W216–W221.

4. Kuleshov, M.V., Jones, M.R., Rouillard, A.D., Fernandez, N.F., Duan, Q., Wang, Z., Koplev, S., Jenkins, S.L., Jagodnik, K.M., Lachmann, A. et al. (2016) Enrichr: a comprehensive gene set enrichment analysis web server 2016 update. Nucleic Acids Research, 44, W90–W97.

5. Davies, E.L., Bell, J.S. and Bhattacharya, S. (2016) Preeclampsia and preterm delivery: A population-based case–control study. Hypertension in Pregnancy, 35, 510–519.

6. Albers, R.E., Kaufman, M.R., Natale, B.V., Keoni, C., Kulkarni-Datar, K., Min, S., Williams, C.R., Natale, D.R.C. and Brown, T.L. (2019) Trophoblast-Specific Expression of Hif-1α Results in Preeclampsia-Like Symptoms and Fetal Growth Restriction. Scientific Reports, 9, 2742.

7. Lee, G., Kil, G., Kwon, J., Kim, S., Yoo, J. and Shin, J. (2009) Oncostatin M as a target biological molecule of preeclampsia. Journal of Obstetrics and Gynaecology Research, 35, 869–875.

8. Wang, X., Zhang, Z., Zeng, X., Wang, J., Zhang, L., Song, W. and Shi, Y. (2018) Wnt/β-catenin signaling pathway in severe preeclampsia. Journal of Molecular Histology, 49, 317–327.

9. Langfelder, P. and Horvath, S. (2007) Eigengene networks for studying the relationships between co-expression modules. BMC Systems Biology, 1, 54.

10. Langfelder, P., Mischel, P.S. and Horvath, S. (2013) When Is Hub Gene Selection Better than Standard Meta-Analysis? PLOS ONE, 8, e61505.

11. Dougan, M., Dranoff, G. and Dougan, S.K. (2019) GM-CSF, IL-3, and IL-5 Family of Cytokines: Regulators of Inflammation. Immunity, 50, 796–811.

12. Chang, S.H. and Dong, C. (2009) IL-17F: Regulation, signaling and function in inflammation. Cytokine, 46, 7–11.

13. Le Mercier, A., Bonnavion, R., Yu, W., Alnouri, M.W., Ramas, S., Zhang, Y., Jäger, Y., Roquid, K.A., Jeong, H.-W., Sivaraj, K.K. et al. (2021) GPR182 is an endothelium-specific atypical chemokine receptor that maintains hematopoietic stem cell homeostasis. Proceedings of the National Academy of Sciences, 118, e2021596118.

14. Bernucci, L., Henríquez, M., Díaz, P. and Riquelme, G. (2006) Diverse Calcium Channel Types are Present in the Human Placental Syncytiotrophoblast Basal Membrane. Placenta, 27, 1082–1095.

15. Zhao, Y., Pasanen, M. and Rysä, J. (2022) Placental ion channels: potential target of chemical exposure. Biology of Reproduction.

16. Moore, K.B., Logan, M.A., Aldiri, I., Roberts, J.M., Steele, M. and Vetter, M.L. (2018) C8orf46 homolog encodes a novel protein Vexin that is required for neurogenesis in Xenopus laevis. Developmental Biology, 437, 27–40.

17. Nadeem, L., Brkic, J., Chen, Y.F., Bui, T., Munir, S. and Peng, C. (2013) Cytoplasmic mislocalization of p27 and CDK2 mediates the anti-migratory and anti-proliferative effects of Nodal in human trophoblast cells. Journal of Cell Science, 126, 445–453.

18. Kaufmann, P., Black, S. and Huppertz, B. (2003) Endovascular Trophoblast Invasion: Implications for the Pathogenesis of Intrauterine Growth Retardation and Preeclampsia. Biology of Reproduction, 69, 1–7.

19. Nakamura, K., Tan, F., Li, Z. and Thiele, C.J. (2011) NGF activation of TrkA induces vascular endothelial growth factor expression via induction of hypoxia-inducible factor-1α. Molecular and Cellular Neuroscience, 46, 498–506.

20. Ding, Z., Zu, S. and Gu, J. (2016) Evaluating the molecule-based prediction of clinical drug responses in cancer. Bioinformatics, 32, 2891–2895.

21. Liu, X., Yin, Z., Xu, L., Liu, H., Jiang, L., Liu, S. and Sun, X. (2021) Upregulation of LINC01426 promotes the progression and stemness in lung adenocarcinoma by enhancing the level of SHH protein to activate the hedgehog pathway. Cell Death & Disease, 12, 173.

22. Giroux-Leprieur, E., Costantini, A., Ding, V.W. and He, B. (2018) Hedgehog Signaling in Lung Cancer: From Oncogenesis to Cancer Treatment Resistance. International Journal of Molecular Sciences, 19, 2835.

23. Bon, E., Driffort, V., Gradek, F., Martinez-Caceres, C., Anchelin, M., Pelegrin, P., Cayuela, M.-L., Marionneau-Lambot, S., Oullier, T., Guibon, R. et al. (2016) SCN4B acts as a metastasis-suppressor gene preventing hyperactivation of cell migration in breast cancer. Nature Communications, 7, 13648.

24. Zhang, C., Guo, H., Li, B., Sui, C., Zhang, Y., Xia, X., Qin, Y., Ye, L., Xie, F.a., Wang, H. et al. (2015) Effects of Slit3 silencing on the invasive ability of lung carcinoma A549 cells. Oncol Rep, 34, 952–960.

25. Yang, C., Yuan, H., Gu, J., Xu, D., Wang, M., Qiao, J., Yang, X., Zhang, J., Yao, M., Gu, J. et al.(2021) ABCA8-mediated efflux of taurocholic acid contributes to gemcitabine insensitivity in human pancreatic cancer via the S1PR2-ERK pathway. Cell Death Discovery, 7, 6.

26. Lagrange, J., Lecompte, T., Knopp, T., Lacolley, P. and Regnault, V. (2022) Alpha-2-macroglobulin in hemostasis and thrombosis: An underestimated old double-edged sword. Journal of Thrombosis and Haemostasis, 20, 806–815.

27. Colaprico, A., Silva, T.C., Olsen, C., Garofano, L., Cava, C., Garolini, D., Sabedot, T.S., Malta, T.M., Pagnotta, S.M., Castiglioni, I. et al. (2016) TCGAbiolinks: an R/Bioconductor package for integrative analysis of TCGA data. Nucleic Acids Res, 44, e71.

28. Sekulic, N., Bassett, E.A., Rogers, D.J. and Black, B.E. (2010) The structure of (CENP-A–H4)2 reveals physical features that mark centromeres. Nature, 467, 347–351.

29. Graser, S., Stierhof, Y.-D. and Nigg, E.A. (2007) Cep68 and Cep215 (Cdk5rap2) are required for centrosome cohesion. Journal of Cell Science, 120, 4321–4331.

30. Bergmann, J.H., Rodríguez, M.G., Martins, N.M.C., Kimura, H., Kelly, D.A., Masumoto, H., Larionov, V., Jansen, L.E.T. and Earnshaw, W.C. (2011) Epigenetic engineering shows H3K4me2 is required for HJURP targeting and CENP-A assembly on a synthetic human kinetochore. The EMBO Journal, 30, 328–340.

31. Stimpson, K.M. and Sullivan, B.A. (2011) Histone H3K4 methylation keeps centromeres open for business. The EMBO Journal, 30, 233–234.

32. Ibrahim, K., Abdul Murad, N.A., Harun, R., Jamal, R., Ibrahim, K., Abdul Murad, N.A., Harun, R., Jamal, R., Ibrahim, K., Abdul Murad, N.A. et al. (2020) Knockdown of Tousled⍰like kinase 1 inhibits survival of glioblastoma multiforme cells. Int J Mol Med, 46, 685–699.

33. Juanes, M.A. (2020) Cytoskeletal Control and Wnt Signaling—APC’s Dual Contributions in Stem Cell Division and Colorectal Cancer. Cancers, 12, 3811.

34. May-Simera, H.L. and Kelley, M.W. (2012) Cilia, Wnt signaling, and the cytoskeleton. Cilia, 1, 7.

35. Ferguson, K.K., Chen, Y.-H., VanderWeele, T.J., McElrath, T.F., Meeker, J.D. and Mukherjee, B. (2017) Mediation of the Relationship between Maternal Phthalate Exposure and Preterm Birth by Oxidative Stress with Repeated Measurements across Pregnancy. Environmental Health Perspectives, 125, 488–494.

36. VanderWeele, T.J. (2016) Mediation Analysis: A Practitioner’s Guide. Annual Review of Public Health, 37, 17–32.

37. VanderWeele, T.J. and Vansteelandt, S. (2010) Odds Ratios for Mediation Analysis for a Dichotomous Outcome. American Journal of Epidemiology, 172, 1339–1348.

38. Ayob, A.Z. and Ramasamy, T.S. (2018) Cancer stem cells as key drivers of tumour progression. Journal of Biomedical Science, 25, 20.

