## Supplementary Data for "*eOmics*: an R package for improved omics data analysis"

### Supplementary Figures

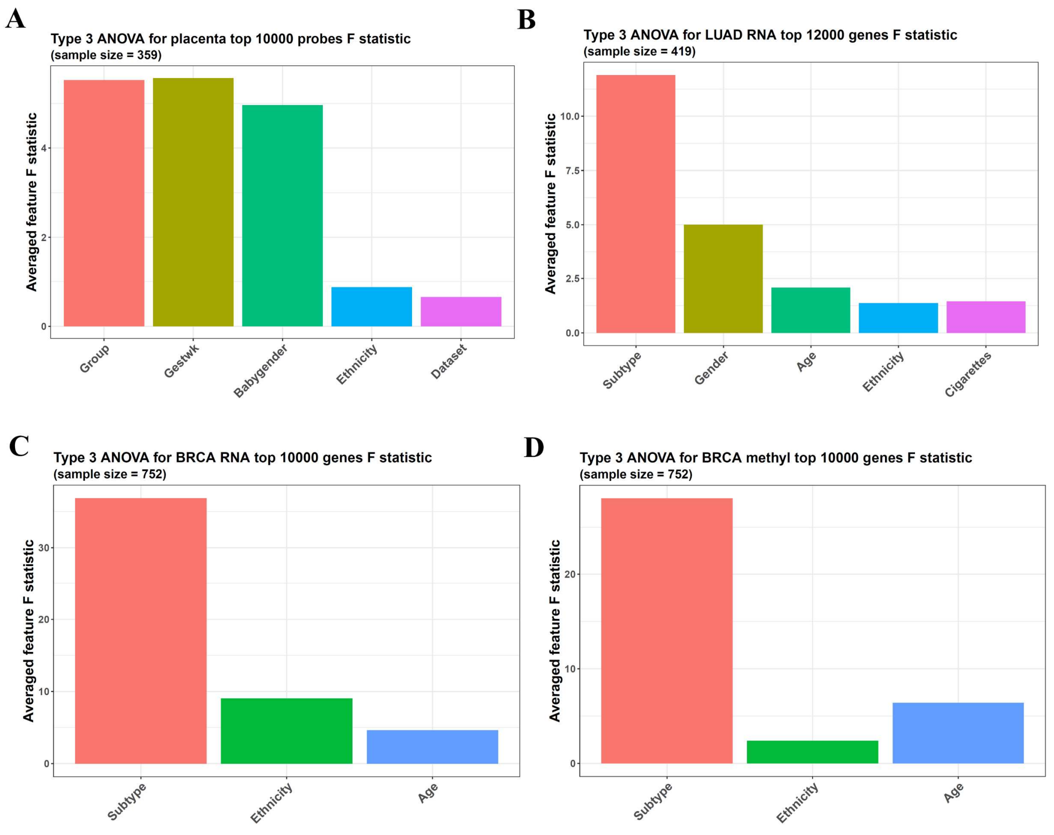

**Figure S1. The ANOVA results of the function *featuresampling*.** (A) For each phenotypic variable shown by the x-axis, *featuresampling* calculates its averaged F statistic across the top 10000 most variable DNAm features in the placenta dataset. Because this dataset is collected from various public ones, the variance of batch differences is also checked, and the F statistic of the variable Dataset is < 1, validating the effect of batch correction on the data. (B) For the top 12000 most variable genes in the LUAD RNA dataset, all the phenotypic variables on the x-axis have an F statistic > 1. (C) and (D) For the top 10000 most variable genes in the BRCA RNA and DNAm gene datasets, the BRCA subtype group, ethnicity, and patient age show an F statistic > 1.

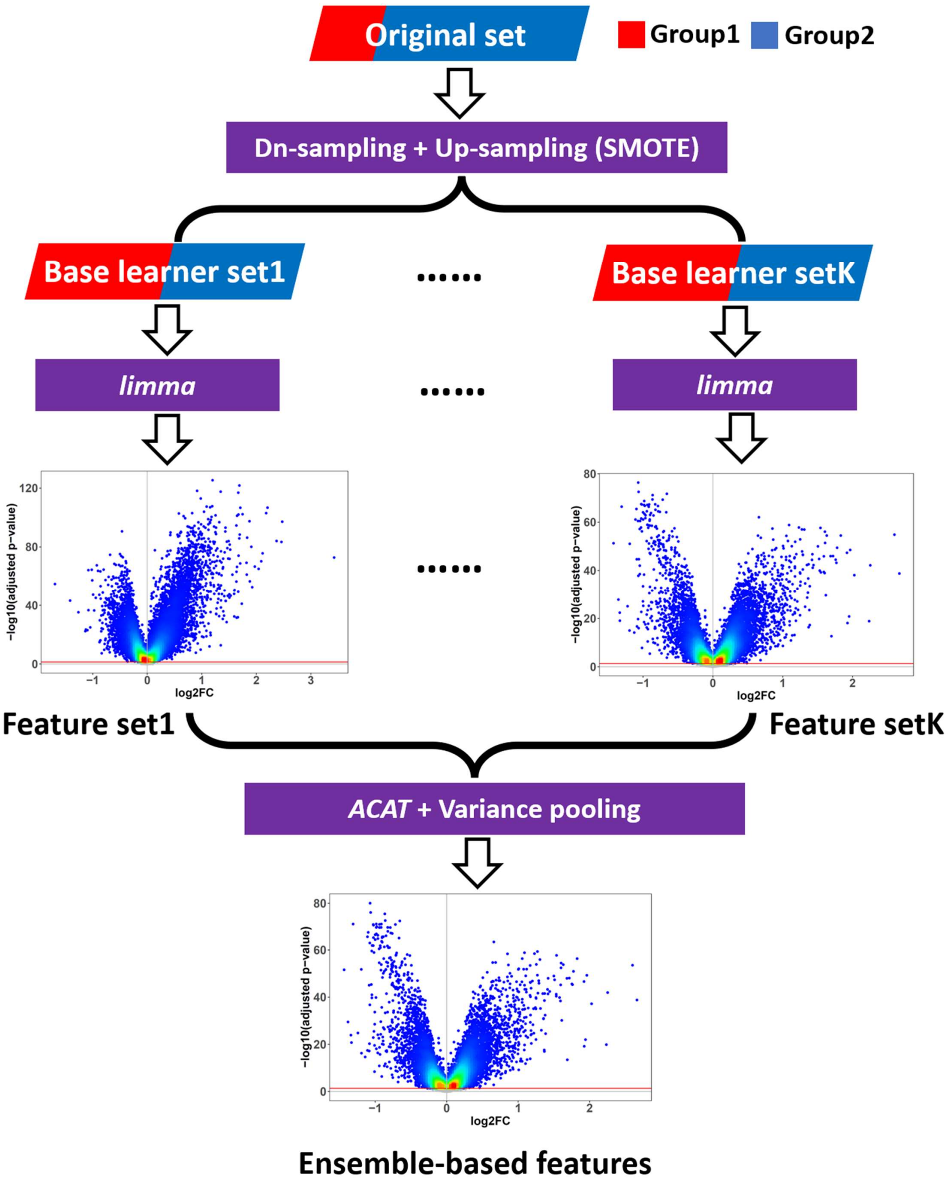

**Figure S2. Ensemble-based *limma* workflow.** The function *difffeatures* performs ensemble-based *limma* with several steps: data distribution adjustment (bagging coupled with SMOTE), *limma* analysis on base learner datasets, and base learner ensemble with *ACAT* (aggregated Cauchy association test) and variance pooling.

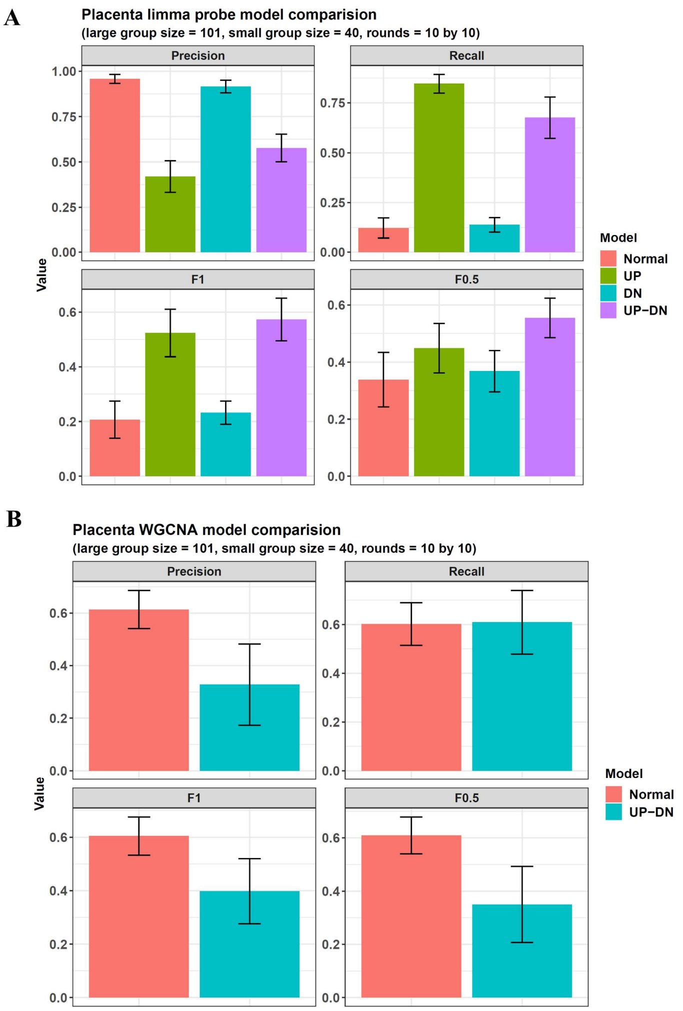

**Figure S3. The ensemble effects on *limma* and *WGCNA* performances.** (A) Simulated experiments show that when ensemble-based *limma* uses up-down sampling to generate base learners, it can call a differential feature set closest to the true one, with higher F1 and F0.5 values than normal *limma* and other methods. (B) However, when combing *WGCNA* with the up-down sampling ensemble framework, the result has more divergence from the true one, i.e., for the gene pairs whose 2 genes are in the same *WGCNA* module, the ones called by the ensemble method are more different from the true pair set, as shown by the F1 and F0.5 statistics.

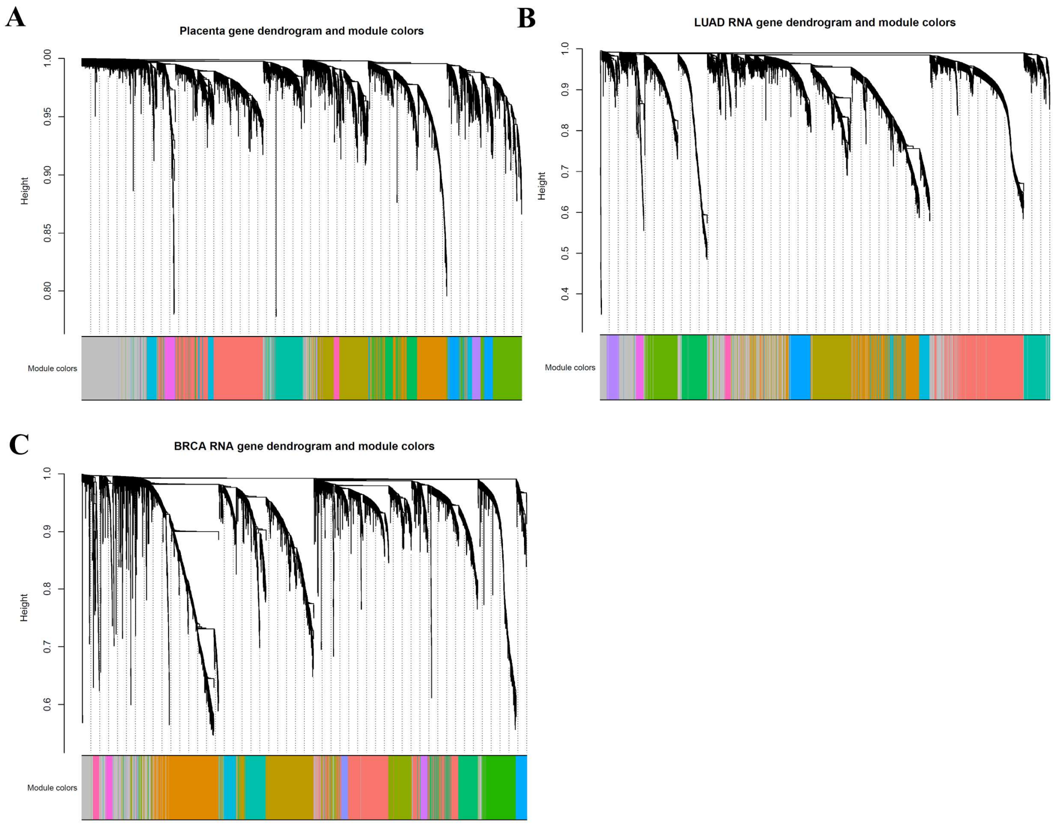

**Figure S4. *WGCNA* dendrograms generated by *diffwgcna*.** The function *diffwgcna* finds 11 modules in the placenta DNAm dataset (A), 11 modules in the LUAD RNA dataset (B), and 13 modules in the BRCA RNA dataset (C).

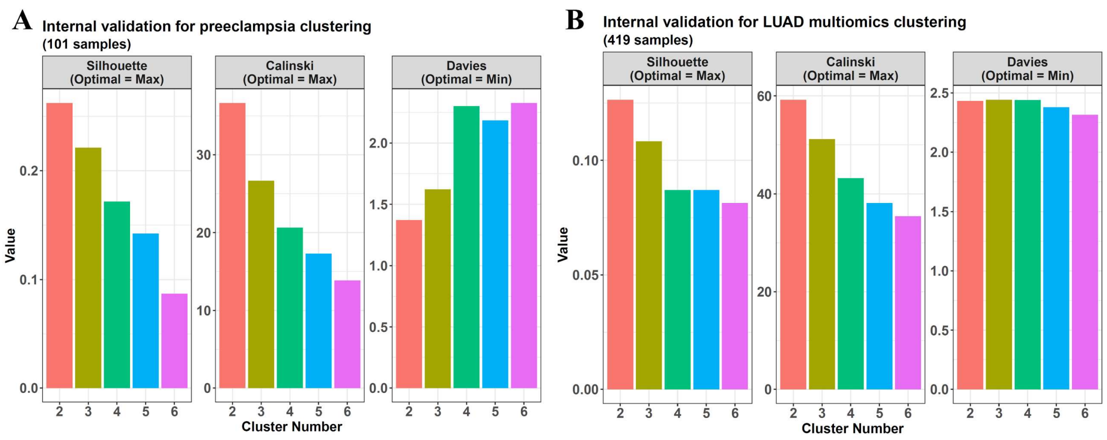

**Figure S5. Internal validation indices for the cluster numbers.** (A) The best cluster number of the preeclampsia DNAm data is 2 with Silhouette index = 0.262, Calinski index = 36.652, and Davies-Bouldin index = 1.371. (B) The best cluster number for the LUAD multi-omics clustering is 2 with Silhouette index = 0.126, Caliniski index = 59.205, and Davies-Bouldin index = 2.432.

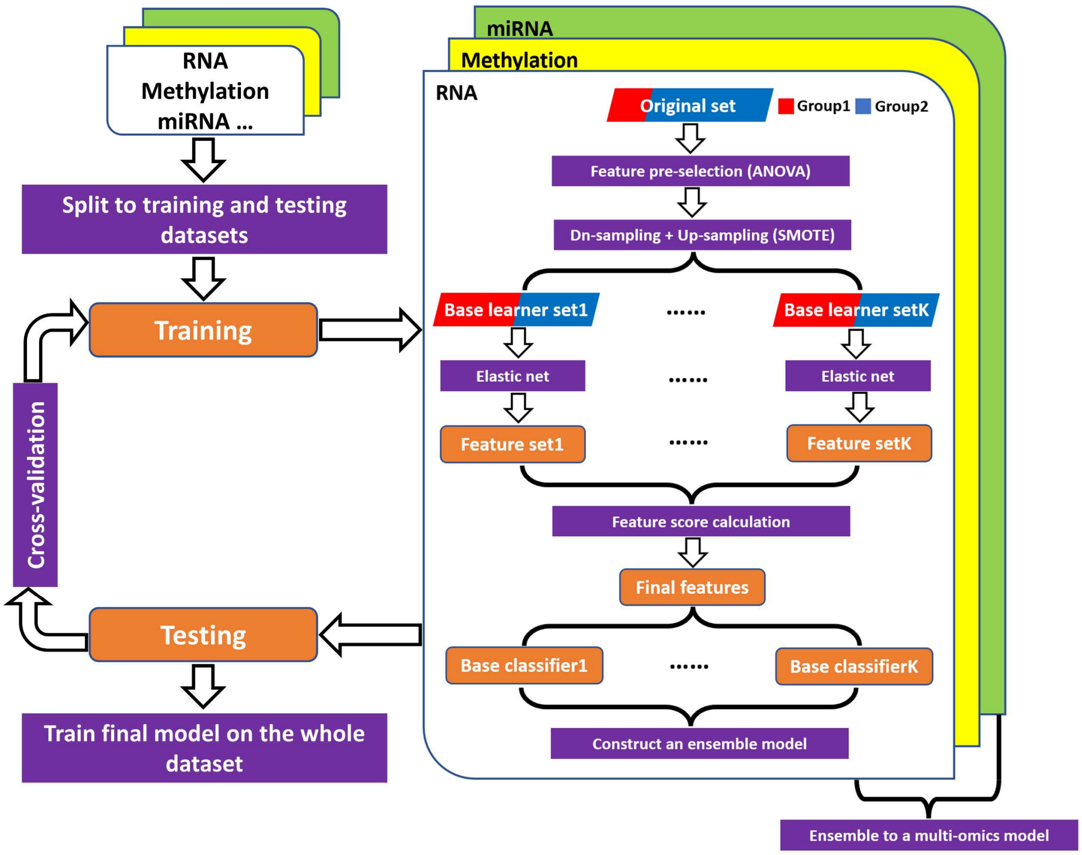

**Figure S6. Workflow of the ensemble-based elastic net model.** The function *omicsclassifier* constructs an ensemble-based elastic net model with several steps: data distribution adjustment (bagging coupled with SMOTE), elastic net base learner training, selected feature integration, multinomial base leaner training, base learner ensemble, and omics model ensemble.

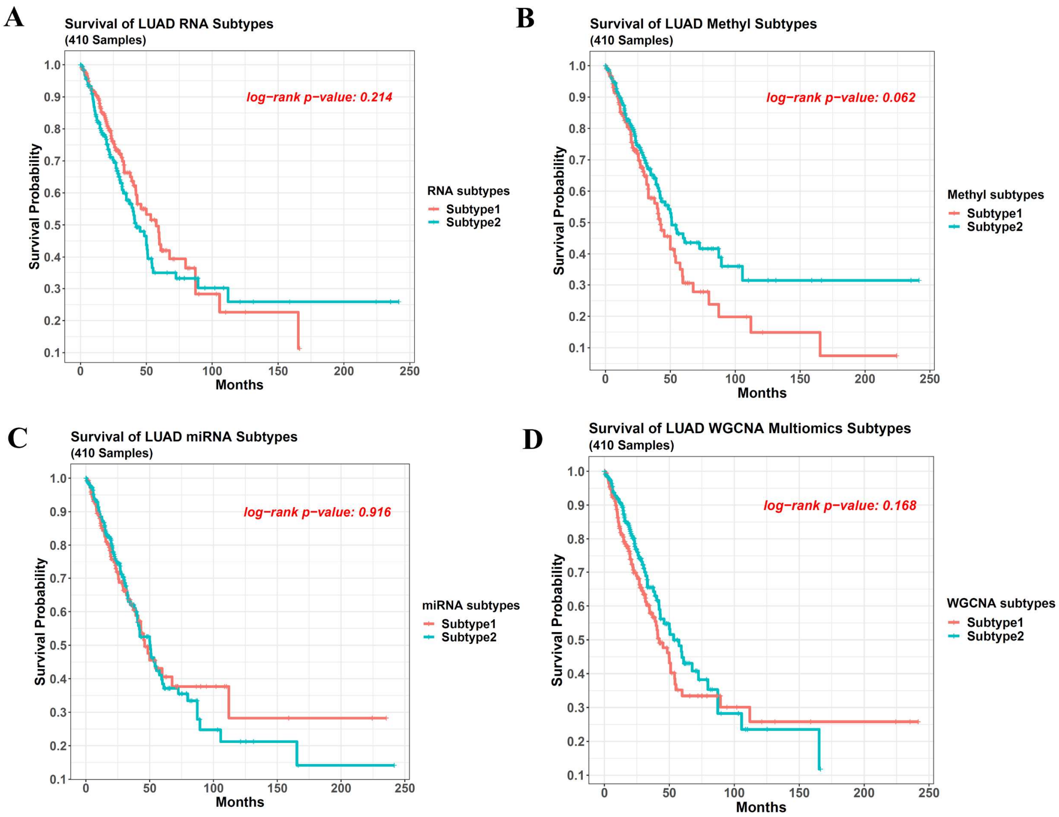

**Figure S7. Survival difference for LUAD subtypes from *multiCCA* single-omic clustering and *wgcnacluster* multi-omics clustering** (A) to (C) The survival difference for *multiCCA* subtypes from RNA, DNAm, and miRNA single-omic data, respectively. (D) That for *wgcnacluster* multi-omics subtypes.

### Supplementary Methods

#### Data collection and preprocessing (*imputemeta*, *sigdmr*, and *probestogenes*)

The Infinium 27K and 450K data on control and preeclampsia human placentas were obtained from 10 GEO datasets: GSE31781 [1], GSE36829, GSE59274 [2], GSE74738 [3], GSE69502 [4], GSE98224 [5, 6], GSE125605, GSE100197, GSE75196, and GSE73375. Then, we used *SeSAMe* to perform data preprocessing and merged the datasets so that only the overlapping probes shared by the Illumina 27K and 450K datasets were kept [7, 8]. The batch difference was adjusted via *ComBat* with GSE98224 data as the reference [9].

GSE98224 also contained RNA expression microarray data for the same placenta samples from Affymetrix Human Gene 1.0 ST Array. The preprocessed RNA expression values were downloaded directly from GEO and used for correlation-based gene function analysis.

The above 10 datasets contained 359 placenta samples (258 control and 101 preeclampsia) and provided the gestational weeks for all of them (from 8 wk to 44.6 wk). In addition, 210 of the 359 samples had baby gender information, and 102 of the 359 samples had ethnicity information. To impute the missing values for baby gender and ethnicity, we used the gender prediction and ethnicity prediction models provided by *SeSAMe*, which predicted them with the methylation beta values. For baby gender, all its missing values could be predicted. Hence, the baby gender information became complete after the prediction, with 210 samples with an original label and 149 with a predicted label. For ethnicity, 257 samples were unlabeled, and 179 could be predicted from their methylation beta values.

Then, the missing ethnicities of the other 78 samples were imputed via the function *imputemeta* in our package. It was based on the *MICE* (multivariate imputations by chained equations) algorithm, which did not impute from beta values, but from other phenotypic variables using Gibbs sampling [10]. After that, the ethnicity information was also complete. Hence, the placenta dataset covered 359 samples. In addition to the methylation data, it contained their phenotypic data on 4 variables: preeclampsia/control group, gestational week, baby gender, and ethnicity.

Among the 10 datasets, 7 were from the 450K platform (except GSE31781, GSE36829, and GSE59274). They were used to call DMRs (DNA methylation regions) significantly related to gestational weeks, which could remove the influence of gestational weeks when performing differential feature calling. The function *sigdmr* in our package conducted this DMR calling based on the R package *bumphunter* [11]. During this process, it used the preeclampsia/control group and baby gender as the confounding factors and performed bootstrapping 100 times to determine the FWERs (family-wide error rates) and p-values for the DMRs. The batch difference among the 7 different 450K datasets was adjusted using *ComBat* in advance, with GSE98224 as the reference.

The RNA, methylation, and miRNA LUAD (lung adenocarcinoma) data were obtained from TCGA, with 419 cancer samples. The RNA and miRNA omics were downloaded as read count tables and converted to log2(TPM + 1). The DNA methylation (DNAm) data were 450K beta values, and the missing values were imputed with the k-nearest neighbors (KNN) method, with specific DNAm probes filtered out, including non-CpG probes, multi-hint probes, SNP probes, and sex chromosome probes. The clinical data were also from TCGA, with missing values imputed by *imputemeta*.

The 752 BRCA (breast invasive carcinoma) cancer samples were from TCGA, and their BRCA

PAM50 subtype information was from the R package *TCGAbiolinks* [12]. Their RNA-seq read counts, 450K DNAm beta values, miRNA-seq read counts, and clinical data were downloaded. Then, the RNA and miRNA data were converted to log2(TPM + 1) values. For the DNAm probe data, after being preprocessed similarly to the LUAD one, they were converted to gene data with our function *probestogenes*, which averaged the probe beta values in gene TSS200, TSS1500, and 1stExon regions as the corresponding gene beta values.

**ANOVA analysis (*featuresampling*)**

The function *featuresampling* in our package was used to perform type-III ANOVA to determine the confounding factors. It constructed a linear regression model to predict each candidate feature with the phenotypic variables. Then, it used type-III ANOVA on the model to calculate the MSS (mean sum of the square), F statistic, and p-value for each phenotypic variable. Finally, each variable got the MSSs, F statistics, and p-values across all the models constructed for all the features, and their means were used to represent the value for the whole dataset.

**Ensemble-based *limma* analysis (*difffeatures* and *sigdmr*)**

The function *difffeatures* in our package performed ensemble-based *limma* analysis following several steps (Figure S2). If its parameter *samplingmethod* were "updn", it would conduct up or down-sampling on each sample group until its final sample number reached the ceiling of $\frac{dataset\ sample\ number}{group\ number}$. Hence, if a group contained too many samples, it would be down-sampled, but if a group only had a few samples, it would be up-sampled.

To implement the up-sampling, *difffeatures* used SMOTE (synthetic minority over-sampling technique) to synthesize new samples via interpolation. If a sample were randomly selected, 1 of its 5 nearest neighbors in the same group would also be sampled. Then, a vector recording the feature value difference between this neighbor and the selected sample would be calculated. Meanwhile, a random number between 0 and 1 would be generated to multiply this vector, and the new vector would be added to the original sample. The result was the synthesized sample with the same group label.

On the other hand, because the large groups lost some samples during down-sampling, a bagging framework was introduced to rescue these samples, so an ensemble model was constructed on the whole dataset. It contained 10 base learner datasets with balanced sample groups. If a base learner did not select a sample during down-sampling, it could still be selected by other base learners, so all the samples could be used for the whole ensemble.

By setting the parameter *samplingmethod* as “updn”, the above method would be called. In addition, if it were set as “up”, only up-sampling would be used to make all the groups reach the largest size; if it were “dn”, all of them would be down-sampled to the smallest size.

After getting the 10 balanced base learner datasets, for each of them, *limma* would be called to find the inter-group differential features. The parameter *confoundings* could be used to assign the confounding factors for the *limma* regression models so that their variances could be removed, which made the differential features only related to sample groups.

In this way, *limma* generated 10 differential feature results from the 10 base learners. Hence, each feature got 10 p-values, which needed to be combined into 1 value to determine its significance. It was a problem similar to the p-value combination in meta-analysis. However, the classic Fisher’s combination for meta-analysis could not be used here because it assumed the p-values independent or weakly dependent, but the p-values here were strongly dependent because the 10 base learner sets were from the same original set. Thus, *difffeatures* used *ACAT* (aggregated Cauchy association test) to perform the p-value combination. Its test statistic was the weighted sum of the Cauchy transformation of p-values and could be approximated by a Cauchy distribution under arbitrary dependency structures [13]. After this combination, each feature got 1 combined p-value, and the Benjamini-Hochberg method was used across all the features to generate their adjusted p-values.

In addition to the 10 p-values from the base learners, each feature had 10 log2FC (fold change) values that needed to be combined. This was performed with the variance pooling approach, which combined them via calculating their weighted sum. The weight for each log2FC was inversely proportional to its base learner's log2FC SEM (standard error of the mean) across all the features [14].

Finally, the inter-group differential features could be identified via their combined p-values, adjusted p-values, and combined log2FCs.

In addition, the function provided another parameter, *removereg*. It accepted a Granges object or a data frame recording the genomic regions related to any confounding factors, and if a differential feature were located in these regions, it would be removed to further avoid influence from the confoundings. For DNAm data, the function *sigdmr* in our package could call the DMRs related to confounding factors, which could be transferred to *removereg* to perform this final feature filtering.

If the parameter *plot* were set as TRUE, *difffeatures* would generate the volcano plot for the features, and if the parameter *balanceadj* were set as FALSE, the function would perform normal *limma* analysis without the sampling and ensemble process.

**Balancing model simulated experiments (*difffeatures*)**

Several simulated datasets were generated from the placenta DNA methylation dataset. First, its small group ($size = x_1$) was fixed as group1. Then, $x_1$ samples were sampled from its large group ($size = x_2 > x_1$) to generate group2. Hence, the result dataset contained 2 groups with the same sample size of $x_1$. The sampling for group2 was performed 10 times so that 10 balanced datasets were generated, each containing the same group1 but a different group2 sampled from the original large group.

For each of the 10 balanced datasets, another 10 rounds of sampling were performed. This time the $x_1$ group2 samples were fixed, and $x_1^2/x_2$ samples were sampled from group1. Hence, 10 imbalanced datasets were from 1 balanced dataset, and a total of 100 imbalanced ones were generated.

After preparing these datasets, we used *limma* and *WGCNA* on the balanced ones and defined their results as true, i.e., *limma* called the true differential features, and *WGCNA* called the true gene pairs with the genes in the same module. Then, we performed ensemble-based *limma* and *WGCNA* for each imbalanced dataset. We tried different sampling methods for their ensembles (up sampling, down sampling, up-down sampling, and no sampling), and we calculated the averaged precision, recall, F1, and F0.5 metrics across the results of the 100 imbalanced datasets so that we could find which sampling generated the differential features or *WGCNA* gene pairs closest to the true ones.

The precision, recall, F1, and F0.5 metrics were calculated as: $precision = \frac{|true\ features\ called|}{|features\ called|}$, $recall = \frac{|true\ features\ called|}{|true\ features|}$, $F1 = \frac{2*precision*recall}{precision+rec}$, $F0.5 = \frac{(1+0.5^2)*precision*recall}{(0.5^2*precision)+recall}$.

**Correlation-based gene functional enrichment (*corenrich*)**

In the placenta data study, for the differential methylation probes called by *difffeatures*, their functional enrichment analysis was conducted with the function *corenrich*. First, it used the paired RNA and DNA methylation data in GSE98224 to calculate the correlation between the differential methylation probes and the gene expression values. Then, if a gene had a PCC (Pearson correlation coefficient) < -0.7 with any hypomethylated site or a PCC > 0.7 with any hypermethylated site, *corenrich* would identify it as an up-regulated gene and *vice versa*. The following enrichment analysis was performed on these RNA genes. Because of the paired RNA-methylation data, only genes with an expression clearly correlated to the changed methylation probes were used. This brought an advantage over the traditional analysis directly on hyper and hypomethylated genes, which assumed gene expression had an anti-correlation with gene methylation in advance, and the functions of hypermethylated genes were annotated as suppressed and *vice versa*. However, many genes did not follow this rule. Even for those with such an anti-correlation, most PCCs were only around -0.5 [15, 16]. Thus, the analysis directly on hyper and hypomethylated genes could not accurately reflect gene expression and function changes. In contrast, *corenrich* conducted the correlation-based gene selection and only focused on the significantly correlated genes, making the analysis more reliable. Then, *corenrich* called the package *enrichR* to find the enriched functions of the up and down-regulated genes.

**Mediation-coupled *WGCNA* analysis (*diffwgcna*)**

The function *diffwgcna* performed mediation-coupled *WGCNA* analysis. It first calculated the *WGCNA* network and modules on the data. Then, it compared the module eigengenes between groups by *limma*, identifying the modules related to the group difference. The use of *limma* to find the differential eigengenes avoided the confounding factors' influence because *limma* regression removed their variance, which was ignored by the traditional *WGCNA* pipeline linking the modules and the phenotype using a correlation coefficient. Furthermore, within each module, *limma* could also be used on its nodes (features) to find the features related to the sample group difference. At these steps, *diffwgcna* also used the parameter *balanceadj* to control whether the ensemble-based *limma* or the normal one should be used.

The *limma* regression, and the traditional correlation method, connected the modules and features with the sample groups. However, this connection was undirected, ignoring the potential causal and mediation relationships among the groups, module features, and modules. Hence, *diffwgcna* included a mediation analysis step to find any directed connections, which could be triggered by setting the parameter *mediation* as TRUE. The mediation models were constructed with the product method [17-19].

Let $Y$ denote an outcome, $A$ an exposure of interest, $M$ a potential mediator, and $C$ a set of baseline covariates, and the corresponding lower-case letters represent realizations of these random variables.

In the case that $Y$ and $M$ were continuous, *diffwgcna* would construct a linear model using the exposure $A$, the mediator $M$, and the covariates $C$ to fit the outcome $Y$ as $E[Y|a,m,c] = \theta_0 + \theta_1 a + \theta_2 m + \theta_3' c$, in which the coefficient $\theta_1$ was interpreted as the direct effect of $A$ on $Y$ ($NDE$, natural direct effect). Also, a second model would be constructed to regress the mediator on the exposure and the covariates as $E[M|a,c] = \beta_0 + \beta_1 a + \beta_2' c$. Then, the product of $\beta_1$ and $\theta_2$ was considered as the indirect effect of $A$ on $Y$

($NIE$, natural indirect effect), which was the effect of the exposure on the mediator times that of the mediator on the outcome. Hence, $NDE = \theta_1$ and $NIE = \theta_2\beta_1$.

On the other hand, if $M$ were continuous but $Y$ were binary, a logistic regression model would be constructed for the outcome as $logit[P(Y = 1|a, m, c)] = \theta_0 + \theta_1 a + \theta_2 m + \theta_3' c$, and the mediator model would still be the linear one as $E[M|a, c] = \beta_0 + \beta_1 a + \beta_2' c$. Then, $NDE$ and $NIE$ would still be $NDE = \theta_1$ and $NIE = \theta_2\beta_1$, but they were on an odds ratio scale.

After that, in the assumption that $M$ mediated the effect of $A$ on $Y$, the proportion mediated ($PCT$) was calculated with $NDE$ and $NIE$. In detail, if $NDE$ and $NIE$ had the same sign, then $PCT = NIE/(NDE + NIE)$. However, if $NDE$ and $NIE$ had opposite signs, which meant there was a suppression mediation effect, then $PCT = |NIE/NDE|$. The 95% confidential intervals (CIs) of $NDE$, $NIE$, and $PCT$ were estimated via bootstrapping 100 times. Accordingly, if the upper and lower confidence limits of $NIE$ had the same sign, a mediation effect would be concluded. Furthermore, if the upper and lower confidence limits of $NDE$ also had the same sign, but $NDE$ and $NIE$ had opposite signs, the mediation would be annotated as a suppression effect.

In addition, IPW (inverse probability weighting) was used to inflate the weights for under-represented observations. If exposure $A$ were binary, covariates $C$ would be used to predict it with logistic regression, and the fitted values for the samples would be the propensity scores to calculate inverse probability weights. A sample with the exposure $A = a$ would get a weight to be used in the mediator model as $\frac{a}{propensity} + \frac{1-a}{1-propensity}$ [20].

On the other hand, if $A$ were continuous, its probability distribution would be estimated as a

Gaussian one of $f_A(A; \mu_1, \sigma_1^2)$, where $\mu_1$ and $\sigma_1^2$ were $A$'s mean and variance across the dataset. At the same time, a linear model would be used to predict $A$ with covariates $C$, generating the probability distribution for each sample as $f_{A|C}(A|C = c; \mu_2, \sigma_2^2)$, where $\mu_2$ and $\sigma_2^2$ were the predictions and residual variance of the model. Then, for a sample with $A = a$ and $C = c$, its weight would be $\frac{f_A(A=a;\mu_1,\sigma_1^2)}{f_{A|C}(A=a|C=c;\mu_2,\sigma_2^2)}$ [21].

The function *diffwgcna* performed the mediation analysis after identifying the sample group's relevant features within each module, and for each of these features, it constructed 2 models. The first one tested the mediation relationship of "module→module feature→group" and the second tested the opposite direction of "group→module feature→module". Hence, both used the feature as the potential mediator $M$ (continuous variable), and they also had the same covariates $C$. The difference was that the first one used the feature's module as the exposure $A$ (continuous module eigengene values) and used the sample group as the outcome $Y$ (binary variable), but the second one swapped them. Finally, the features mediating their module's effect on the group, or *vice versa*, were identified by the 95% CIs of the 2 models. Sometimes a feature was significant in both models and would be discarded.

If the parameter *balanceadj* were FALSE, this mediation analysis would be performed on the modules and features of the original dataset. However, if this parameter were TRUE, it would be performed on the modules and features of each base learner dataset first, and then the results would be ensembled. For $NDE$, the mean of all the base leaner $NDEs$ would be the aggregated one, and its upper confidence limit would be $\overline{NDE} + \sqrt{\frac{\sum_{i=1}^{n}(\widehat{NDE_i} - NDE_i)^2}{n}}$, where $\overline{NDE}$ was the mean of all the base learner $NDEs$, $NDE_i$ was the $NDE$ of the $i$th base learner, and $\widehat{NDE_i}$ was its upper confidence limit. The lower confidence limit would be calculated similarly, and the $NIE$ and $CPT$ results would also be aggregated in this way.

**Multi-omics clustering (*multiCCA* and *wgcnacluster*)**

The function *multiCCA* performed multi-omics clustering. It first scaled the omics data so all the features had a unified mean of 0 and a standard deviation of 1. If only 2 omics were used for clustering, the covariance matrix between them would be calculated, and CCA (canonical correlation analysis) would be used on it, generating the top 30 CC components for each omic. Hence, in each single-omic dataset, all the samples got 30 new features (the CCs), and a matrix was formed. Its row number was the sample number, and its column number was the CC number. In total, 2 such matrices were generated for the 2 omics. At the same time, a weight was calculated for each matrix, determined by the variance it explained for its single-omic data. Next, the weights of the 2 matrices were scaled to the sum of 1, and their weighted sum was calculated, generating a merged matrix. If there were a third omic, this merged matrix would be treated as a single omic, and the third omic was the other, then the same process was used to merge them, and if there were also the fourth or other omics, this process would be repeated to combine all of them into one merged matrix, which would undergo $k$-means clustering.

To determine the optimal $k$ value for the $k$-means clustering, *multiCCA* accepted multiple candidate $k$ values. Then, for each of them, 3 internal validation indices were calculated after the clustering. They were: 1) the Silhouette index as $Silhouette = \frac{1}{k}\sum_{i=1}^{k}\left\{\frac{1}{|C_i|}\sum_{x\in C_i}\frac{b(x)-a(x)}{\max[b(x),a(x)]}\right\}$, where $a(x) = \frac{1}{|C_i|-1}\sum_{y\in C_i, x\neq y} d(x,y)$, $b(x) = min_{j,j\neq i}\left[\frac{1}{|C_j|}\sum_{y\in C_j} d(x,y)\right]$, 2) the Calinski index as $Calinski = \frac{\sum_i^k |C_i| d^2(c_i,c)/(n-1)}{\sum_i^k \sum_{x\in C_i} d^2(x,c_i)/(n-k)}$, and 3) the Davies-Bouldin index as $Davis - Bouldin = \frac{1}{k}\sum_i^k max_{j,j\neq i}\left\{\left[\frac{1}{|C_i|}\sum_{x\in C_i} d(x,c_i) + \frac{1}{|C_j|}\sum_{x\in C_j} d(x,c_j)\right]/d(c_i,c_j)\right\}$, where $d(x,y)$ was the distance between samples $x$ and $y$

on the merged matrix, $C_i$ was the $i$th cluster, $c_i$ was its center, $c$ was the center of the whole data. The optimal $k$ value was the one with the largest Silhouette and Calinski indices and the smallest Davis-Bouldin index.

Besides, *multiCCA* could perform consensus clustering by bootstrapping the features 100 times. Each time, a new multi-omics dataset was generated, and the above CCA-based clustering was used. Finally, the 100 clustering results would be aggregated to get the consensus.

If the dataset was not multi-omics but single-omic, *multiCCA* could also perform the above process to cluster, but the CCA would be changed to PCA on the covariance matrix of the single-omic dataset.

In addition, our package contained another function, *wgcnacluster*, which could also perform multi-omics clustering and had almost the same steps as *multiCCA*. However, its method to generate the merged matrix before *k*-means clustering was different. For each omic, *wgcnacluster* did not use CCA but used *WGCNA* to generate the *WGCNA* module eigengenes for its samples. Then, the ones from different omics were directly combined, generating a final matrix containing multi-omics *WGCNA* module eigengenes. After that, *k*-means would be used on it to perform the clustering.

**Multi-omics classification (*omicsclassifier* and *pairedensemblepredict*)**

In addition to clustering, multi-omics classification could also be performed with our package, depending on its function *omicsclassifier*. It scaled the omics data first, making all the features have a unified mean of 0 and a standard deviation of 1. Then, it performed type-III ANOVA on each feature to screen the ones with a p-value < 0.05 to the response variable (sample classes), and they would be used for the classification following several steps (Figure S6).

For each single-omic dataset, if the parameter *balanceadj* were set as TRUE, *omicsclassifier* would perform sampling to generate 10 balanced base learner datasets, similar to the function *difffeatures*, balancing the sample group sizes in the base learner datasets. Then, each dataset would be used to train a base learner via elastic net regularization, $\min_{(\beta_{0k},\beta_k)\in\mathbb{R}^{p+1}} \left[\frac{1}{N}\sum_{i=1}^{N}(-\sum_{k=1}^{K} y_{il}\,(\beta_{0k} + x_i^T\beta_k) + \log\,(\sum_{l=1}^{K} e^{\beta_{0l}+x_i^T\beta_l}))\right] + \lambda\left[(1-\alpha)\,\frac{1}{2}\parallel \beta \parallel_F^2 + \alpha\sum_{j=1}^{p}\parallel \beta_j \parallel_1\right]$ [22, 23]. In the formula, the first part was the multinomial negative log-likelihood, and the second was the multiclass elastic net penalty. The $\alpha$ parameter was set as 0.5, which controlled the balance of L1 and L2 penalties. At the same time, the regularization constant $\lambda$ was chosen during a 10-fold cross-validation. The function chose the $\alpha$-$\lambda$ combination giving the minimum cross-validation error, and used it to construct the elastic net model.

Each base learner used this elastic net method to select a set of features. Then, their features were combined, and those with the top scores were selected, which were calculated referring to a previous method predicting drug responses [24]. It used the formula, $\left(\mathcal{F}_p^+ - \mathcal{F}_p^-\right) * \overline{\beta_p}$, where $\mathcal{F}_p^+ = \frac{1}{K}\sum_{k=1}^{K} I\left(\beta_p^{(k)} > 0\right)$ and $\mathcal{F}_p^- = \frac{1}{K}\sum_{k=1}^{K} I\left(\beta_p^{(k)} < 0\right)$ represented the percentages of base learners with a coefficient larger or smaller than 0 for the *p*th feature selected for a specific sample class, and $\overline{\beta_p} = \frac{1}{K}\sum_{k=1}^{K}\beta_p^{(k)}$ was the averaged coefficient value of this feature across all the *K* base learners.

Then, the selected features were returned to each base learner dataset to generate a multinomial model. Finally, these new base learners were ensembled together, which was achieved by assigning each of them a weight as $0.5 * \log\frac{ACC}{1-ACC}$, where $ACC$ was the base learner prediction accuracy on the original training dataset. The predicted class distribution of the whole ensemble was the weighted sum of the base learner ones. Only the base learners with an $ACC$ greater than 0.5 would be included in the ensemble, and the base learner weights would be scaled to the sum of 1.

If the dataset transferred to *omicsclassifier* were single-omic, the above process would return the final classification result, but if it were multi-omics, it would be used on each single-omic first, and their predicted class distributions would be further aggregated with the weighted sum method above. However, the weights were not calculated from each base learner but from each single-omic model accuracy.

In addition, *omicsclassifier* also contained a parameter *nfold*, which could assign a fold number to perform cross-validation training. If it were NULL, this step would be skipped, and only the classifier trained from the whole data would be returned. It could be further transferred to another function, *pairedensemblepredict*, which predicted new sample classes with this classifier.

**Structure-based gene functional enrichment (*diffwgcna* and *topoenrich*)**

The *WGCNA* modules' functional enrichment could be performed by *diffwgcna*. Unlike traditional enrichment methods that only covered a module's nodes (genes), *diffwgcna* further considered its network structure, reflected by the network edges and their weights.

In our edge-based analysis, the function terms of each edge (gene-gene pair) were defined as the intersection of its 2 genes'. Then, 2 methods could be used on them to perform enrichment: 1) edge weight shuffling and 2) hypergeometric test.

For the edge weight shuffling method, *diffwgcna* shuffled a module's edge weights 1000 times, and each time, all edges with a specific function term were summed up. Hence, 1000 weight sums were generated for this term to form a distribution. Then, its original sum in the module was mapped to it to get the probability of a greater sum, which was the p-value. This method considered all gene pairs in the module, so even if there were no edges between 2 nodes, this gene pair would still be shuffled but with a weight of 0.

For the other hypergeometric test method, if not considering weights, the test for a function term could be calculated with 4 numbers, including 1) $N$: the total number of edges in the background, 2) $n$: that in the module, 3) $M$: the number of edges with that function term in the background, and 4) $m$: that in the module. Hence, the final p-value was $P(X > k) = \sum_{m=k+1}^{\min(M,n)} \frac{\binom{M}{m}\binom{N-M}{n-m}}{\binom{N}{n}}$, where $k$ was the realization of $m$ in the module.

If further considering edge weights in the enrichment, the hypergeometric test would become difficult because it needed to calculate combinations, such as $\binom{N}{n} = \frac{N!}{(N-n)!n!}$, where the factorials required non-negative integers. However, the edge weights were not restricted to integers. Hence, *diffwgcna* referred to a gene-based enrichment tool, *WEAT*, which solved this problem by introducing the $\Gamma$ function [25]. Similarly, *diffwgcna* used it because it was a common extension of factorial to complex numbers as $x! = \Gamma(x+1) = \int_0^{\infty} t^x e^{-t} dt$, where $x$ was a complex number. Accordingly, the hypergeometric test on edge weights became $P(X > k) = \int_k^{\min(M,n)} \frac{\binom{M}{m}\binom{N-M}{n-m}}{\binom{N}{n}} dm$. The factorials for calculating $\binom{N}{n}$ were solved by the $\Gamma$ function, and $N$ and $n$ were the total edge weights in the background and the module. In addition, the edge weights were normalized in advance as $w_i' - \frac{1}{n}\sum_{i=1}^{n} w_i' + 1$, where $w_i' = \frac{w_i - \min(W)}{\max(W) - \min(W)}$, and $W$ was the vector of all edge weights, including the $i$th as $w_i$.

However, because *WGCNA* constructed a fully-connected network across all the input genes, the gene pair number in the background would be huge due to the gene combination, bringing a burden of calculating so many gene pairs. Hence, only the top 1% edges with the highest TOM (topological overlap matrix) weights would be considered as the background, and for a module, only its edges belonging to this top 1% would be in the enrichment analysis.

For both the shuffling and hypergeometric methods, their p-values would be adjusted with Benjamini-Hochberg correction, and if changing the edge weights to gene weights, the methods could also be used on genes.

These structure-based analyses could also be performed with the function *topoenrich* in the package, and besides the *WGCNA* modules, it also analyzed other gene networks. The databases used for the enrichment could be chosen from Reactome, GOBP, GOCC, GOMF, and KEGG.

**Multi-omics regulatory analysis (*lassomediation*)**

The function *lassomediation* performed multi-omics regulatory analysis. Its parameters *rnadat* and *methyldat* accepted RNA and DNAm, or other omics data, respectively. Then, the analysis would follow several steps.

First, if its parameter *calldifffeatures* were set as TRUE, the function would select differential RNA and DNAm features between different sample groups. It generated several balanced base learner datasets from the RNA and DNAm data and then used *limma* on each. After that, *ACAT* combination and variance pooling were used to ensemble the base learner results. It was similar to the process of *difffeatures*, but the samples in the RNA and DNAm base learner sets were generated in the same sampling step to guarantee they had paired RNA and DNAm samples, which was important to the downstream analysis.

Then, an ensemble-based LASSO model would be constructed for each differential RNA gene, where the RNA was the response variable, and all the differential DNAm features were candidate predictors. This model used the base learner datasets above, training a LASSO model for each of them to select a set of DNAm features to fit the RNA. It was similar to the steps in *omicsclassifier*, but the LASSO model here was to perform regression rather than classification. After that, the base leaner DNAm features were combined, using the method in *omicsclassifier*, and the final features would be returned to each base learner dataset to train a linear regression model for its RNA.

Finally, the new base learner results were ensembled by assigning each of them a weight as $0.5 * \log\frac{R^2}{1-R^2}$ with the following scale to the sum of 1, where $R^2$ was the regression R square between the true RNA value and the base learner-predicted one for the original dataset. Only the base learners with an R square > 0.5 would be used for the ensemble, and the final RNA prediction was the weighted sum of the base learners.

After this LASSO feature selection, if the final ensemble predicted the RNA value with an R square > 0.5, its selected DNAm features would go to the mediation analysis. For each of them, two causal directions would be analyzed, i.e., "sample group→RNA→DNAm feature" and "DNAm feature→RNA→sample group". This step was also performed on each base learner dataset, and the final result was their ensemble, which was similar to the meditation tests in *diffwgcna*.

In addition, if the function parameter *balanceadj* were set as FALSE, the LASSO and mediation models could be constructed without the ensemble framework. If the parameter *calldifffeatures* were set as FALSE, the function could skip the step for differential RNA and DNAm feature identification. All the RNA genes transferred to *lassomediation* would go through this pipeline.

### References

1. Novakovic B, Yuen RK, Gordon L et al. Evidence for widespread changes in promoter methylation profile in human placenta in response to increasing gestational age and environmental/stochastic factors, BMC Genomics 2011;12:529.
2. Chu T, Bunce K, Shaw P et al. Comprehensive analysis of preeclampsia-associated DNA methylation in the placenta, PLoS One 2014;9:e107318.
3. Hanna CW, Peñaherrera MS, Saadeh H et al. Pervasive polymorphic imprinted methylation in the human placenta, Genome Res 2016;26:756-767.
4. Price EM, Peñaherrera MS, Portales-Casamar E et al. Profiling placental and fetal DNA methylation in human neural tube defects, Epigenetics Chromatin 2016;9:6.
5. Leavey K, Wilson SL, Bainbridge SA et al. Epigenetic regulation of placental gene expression in transcriptional subtypes of preeclampsia, Clin Epigenetics 2018;10:28.
6. Wilson SL, Leavey K, Cox BJ et al. Mining DNA methylation alterations towards a classification of placental pathologies, Hum Mol Genet 2018;27:135-146.
7. Zhou W, Triche TJ, Jr., Laird PW et al. SeSAMe: reducing artifactual detection of DNA methylation by Infinium BeadChips in genomic deletions, Nucleic Acids Res 2018;46:e123.
8. Triche TJ, Jr., Weisenberger DJ, Van Den Berg D et al. Low-level processing of Illumina Infinium DNA Methylation BeadArrays, Nucleic Acids Res 2013;41:e90.
9. Leek JT, Johnson WE, Parker HS et al. The sva package for removing batch effects and other unwanted variation in high-throughput experiments, Bioinformatics 2012;28:882-883.
10. van Buuren S, Groothuis-Oudshoorn K. mice: Multivariate Imputation by Chained Equations in R, Journal of Statistical Software 2011;45:1 - 67.
11. Jaffe AE, Murakami P, Lee H et al. Bump hunting to identify differentially methylated regions in epigenetic epidemiology studies, International Journal of Epidemiology 2012;41:200-209.
12. Colaprico A, Silva TC, Olsen C et al. TCGAbiolinks: an R/Bioconductor package for integrative analysis of TCGA data, Nucleic Acids Res 2016;44:e71.
13. Liu Y, Xie J. Cauchy Combination Test: A Powerful Test With Analytic p-Value Calculation Under Arbitrary Dependency Structures, Journal of the American Statistical Association 2020;115:393-402.
14. Hodos RA, Strub MD, Ramachandran S et al. Integrative genomic meta-analysis reveals novel molecular insights into cystic fibrosis and ΔF508-CFTR rescue, Scientific Reports 2020;10:20553.
15. Teschendorff AE, Zhu T, Breeze CE et al. EPISCORE: cell type deconvolution of bulk tissue DNA methylomes from single-cell RNA-Seq data, Genome Biol 2020;21:221.
16. Jiao Y, Widschwendter M, Teschendorff AE. A systems-level integrative framework for genome-wide DNA methylation and gene expression data identifies differential gene expression modules under epigenetic control, Bioinformatics 2014;30:2360-2366.
17. Ferguson KK, Chen Y-H, VanderWeele TJ et al. Mediation of the Relationship between Maternal Phthalate Exposure and Preterm Birth by Oxidative Stress with Repeated Measurements across Pregnancy, Environmental Health Perspectives 2017;125:488-494.
18. VanderWeele TJ. Mediation Analysis: A Practitioner's Guide, Annual Review of Public Health 2016;37:17-32.
19. VanderWeele TJ, Vansteelandt S. Odds Ratios for Mediation Analysis for a Dichotomous Outcome, American Journal of Epidemiology 2010;172:1339-1348.
20. Jo B, Stuart EA, MacKinnon DP et al. The Use of Propensity Scores in Mediation Analysis, Multivariate

Behavioral Research 2011;46:425-452.
21. Naimi AI, Moodie EEM, Auger N et al. Constructing Inverse Probability Weights for Continuous Exposures: A Comparison of Methods, Epidemiology 2014;25:292-299.
22. Friedman J, Hastie T, Tibshirani R. Regularization Paths for Generalized Linear Models via Coordinate Descent, J Stat Softw 2010;33:1-22.
23. Tibshirani R, Bien J, Friedman J et al. Strong rules for discarding predictors in lasso-type problems, J R Stat Soc Series B Stat Methodol 2012;74:245-266.
24. Ding Z, Zu S, Gu J. Evaluating the molecule-based prediction of clinical drug responses in cancer, Bioinformatics 2016;32:2891-2895.
25. Fan R, Cui Q. Toward comprehensive functional analysis of gene lists weighted by gene essentiality scores, Bioinformatics 2021;37:4399-4404.
