## Supplementary material for "*eOmics*: an R package for improved omics data analysis": Tutorial

Tutorial for R package eOmics

Yu Liu

2023-01-23

### Introduction

Many computational tools have been developed for high-throughput omics data, and some are very popular, such as *limma* *[1]*, *WGCNA* *[2]*, and *EnrichR* *[3]*. However, they also exhibit disadvantages in some special cases, such as imbalanced data analysis, causal inference, gene network functional enrichment, etc. Hence, we developed the R package *eOmics* (enhanced omics) to provide a comprehensive pipeline with these problems addressed. It combines an ensemble framework with *limma* and *WGCNA*, improving their performance on imbalanced data. Moreover, it couples a mediation model with *WGCNA*, so the causal relationship among *WGCNA* modules, module features, and phenotypes can be found, and this model is also used to explore the relationship between different omics. In addition, our package has some novel functional enrichment methods, capturing the influence of topological structure on gene set functions. Finally, it contains multi-omics clustering and classification functions to facilitate machine-learning tasks. Some basic functions, such as missing value imputation, are also available. This tutorial will introduce the main functions of *eOmics*.

### Package installation

Code of *eOmics* is freely available at <https://github.com/yuabrahamliu/eOmics>.

The following commands can be used to install this R package.

library(devtools)

install_github("yuabrahamliu/eOmics")

### Data preparation

This tutorial will use the data accompanying the *eOmics* package. They contain a matrix covering the DNA methylation (DNAm) data from 359 human placenta samples, which are collected from 10 different GEO datasets based on the platforms of Illumina 27K and 450K. The matrix columns represent the samples, and the rows represent DNAm probes. Totally 18626 probes are in the matrix and are shared high-quality probes among the samples. The probe values are beta values and have been preprocessed with batch difference adjusted. Among the 359 samples, 258 are normal samples, and the remaining 101 are disease samples with preeclampsia pregnancy complications.

In addition to this matrix, a data frame with this package records the metadata for the 359 samples. It includes the sample IDs (column “sampleid”, corresponding to the column names of the beta value matrix), their preeclampsia/control status (column “Group”), the gestational ages (weeks) of the samples (column “Gestwk”), the baby gender (column “Babygender”), the maternal ethnicity (column “Ethnicity”), and the original GEO dataset IDs of the samples (column “Dataset”).

Another data frame this tutorial needs records gestational age-relevant DNA methylation regions (DMRs). Gestational age is a strong confounding factor for preeclampsia analysis. Hence, we called its relevant DMRs from 7 450K datasets (the ones in the above 10 GEO datasets, except the 3 27K ones GSE31781, GSE36829, and GSE59274). We transferred them to the function *sigdmr* in *eOmics* to call gestational age-relevant DMRs, which generated the data frame, with its rows as DMRs, and its columns “seqnames”, “start”, “end”, and “strand” are used to show the genomic coordinates of the DMRs necessary for our next analysis.

Attach *eOmics* to the R session and look at these data.

library(eOmics)

betas <- system.file("extdata", "placentabetas.rds", package = "eOmics")
betas <- readRDS(betas)

pds <- system.file("extdata", "placentapds.rds", package = "eOmics")
pds <- readRDS(pds)

gestdmrs <- system.file("extdata", "gestDMR.rds", package = "eOmics")
gestdmrs <- readRDS(gestdmrs)

The beginning parts of these data are shown below.

```
#The DNAm betas matrix

betas[1:6,1:6]
#>             GSM788417  GSM788419  GSM788420  GSM788421  GSM788414  GSM788415
#> cg00000292 0.65961366 0.67591141 0.65709651 0.66077820 0.66847653 0.67436406
#> cg00002426 0.53516824 0.53883284 0.53683120 0.53990206 0.53804637 0.53543344
#> cg00003994 0.17674229 0.16432771 0.16631494 0.16803355 0.16416337 0.16852233
#> cg00007981 0.03553198 0.02934814 0.03189098 0.02765783 0.02768899 0.02764183
#> cg00008493 0.43619912 0.42998023 0.43761396 0.44973257 0.46281094 0.45935308
#> cg00008713 0.07431056 0.06367040 0.06764089 0.06218142 0.05267028 0.05397539
#The metadata for the samples
head(pds)
#>    sampleid   Group Gestwk Babygender Ethnicity  Dataset
#> 1 GSM788417 Control      8          M     White GSE31781
#> 2 GSM788419 Control      8          M     White GSE31781
#> 3 GSM788420 Control      8          M     White GSE31781
#> 4 GSM788421 Control      9          M     White GSE31781
#> 5 GSM788414 Control     12          F     Asian GSE31781
#> 6 GSM788415 Control     12          M     White GSE31781

table(pds$Group)
#>
#>      Control Preeclampsia
#>          258          101

table(pds$Dataset)
#>
#> GSE100197 GSE125605  GSE31781  GSE36829  GSE59274  GSE69502  GSE73375  GSE74738
#>        65        41        30        48        23        16        36        28
#>  GSE75196  GSE98224
#>        24        48
#The gestational age-relevant DMR data
head(gestdmrs)
#>       seqnames     start       end width strand        value        area
#> DMR_1    chr17  40687519  40687519     1      * -0.008064685 0.008064685
#> DMR_2    chr17  40687596  40687596     1      * -0.006044539 0.006044539
#> DMR_3    chr12 104697193 104697545   353      *  0.002243535 0.015704742
#> DMR_4     chr6  33156836  33157222   387      *  0.002037436 0.018336920
#> DMR_5    chr11  63871000  63871000     1      *  0.003379808 0.003379808
#> DMR_6    chr13  23309774  23309774     1      *  0.003346264 0.003346264
#>       cluster indexStart indexEnd      L clusterL      p.value fwer p.valueArea
#> DMR_1   70224      58553    58553      1       14 4.164931e-05 0.01  0.06772178
#> DMR_2   70224     198622   198622      1       14 1.249479e-04 0.03  0.09683465
#> DMR_3   39798      27573   341167 313595        7 2.498959e-03 0.48  0.02119950
#> DMR_4  134001       7867   314517 306651        9 3.498542e-03 0.61  0.01378592
#> DMR_5   29562     271167   271167      1        9 3.998334e-03 0.51  0.19620991
#> DMR_6   43231     212100   212100      1        8 4.164931e-03 0.52  0.19841733
#>       fwerArea
#> DMR_1     1.00
#> DMR_2     1.00
#> DMR_3     1.00
#> DMR_4     0.94
#> DMR_5     1.00
#> DMR_6     1.00
```

### Confounding factor analysis

We first check the relationship between the methylation beta values in the betas matrix and the phenotypic variables in the pds metadata using the function *featuresampling* in our package. It performs type-III ANOVA to measure the variance of each phenotypic variable account for the beta value dataset.

We only perform the analysis on the top 10000 most variable DNAm probes in the data, which can be selected by *featuresampling*. Its parameter betas accepts our betas placenta data matrix, and the parameter topfeatures can be set as 10000 so that the top 10000 most variable probes will be selected. Another parameter, variancetype, is used to define how to calculate the variance, and we set it as “sd”, meaning that standard deviation will be used. The parameter threads defines the threads number for parallelization.

```
top10k <- featuresampling(betas = betas,
                          topfeatures = 10000,
                          variancetype = "sd",
                          threads = 4)

top10k$betas[1:6,1:6]
#>            GSM788417 GSM788419 GSM788420 GSM788421 GSM788414 GSM788415
#> cg00000292 0.6596137 0.6759114 0.6570965 0.6607782 0.6684765 0.6743641
#> cg00002426 0.5351682 0.5388328 0.5368312 0.5399021 0.5380464 0.5354334
#> cg00003994 0.1767423 0.1643277 0.1663149 0.1680336 0.1641634 0.1685223
#> cg00008493 0.4361991 0.4299802 0.4376140 0.4497326 0.4628109 0.4593531
#> cg00013618 0.7859429 0.7868591 0.7933922 0.7958716 0.8006630 0.8118193
#> cg00014837 0.8522323 0.8525327 0.8516924 0.8546466 0.8610099 0.8546916

dim(top10k$betas)
#> [1] 10000   359

head(top10k$varicanceres)
#>                   SD
#> cg03729431 0.1848922
#> cg16063666 0.1643842
#> cg05790038 0.1456048
#> cg03316864 0.1454261
#> cg11009736 0.1393115
#> cg20322876 0.1348288
```

The result top10k is a list. Its slot betas contains the data matrix only with the top 10000 most variable DNAm probes, and the other slot varicanceres records the standard deviation of the original 18626 probes calculated by *featuresampling*. Then, we further use this function to perform ANOVA on the slot betas. We transfer our metadata pds to its parameter pddat, so all its variables except the first column “sampleid” will be analyzed by ANOVA. The parameter anova should be set as TRUE to make the function perform ANOVA.

The parameter plotannovares is used to indicate whether it is needed to plot the ANOVA result for the data, and featuretype and plottitilesuffix are characters that need to be shown in the plot title and can also be set as NULL. titilesize and textsize are used to control the font sizes of the figure title and texts.

```
anovares <- featuresampling(betas = top10k$betas,
                            pddat = pds,
                            anova = TRUE,

                            plotannovares = TRUE,
                            featuretype = "probe",
                            plottitlesuffix = "placenta",
                            titlesize = 18,
                            textsize = 16,

                            threads = 4)
```

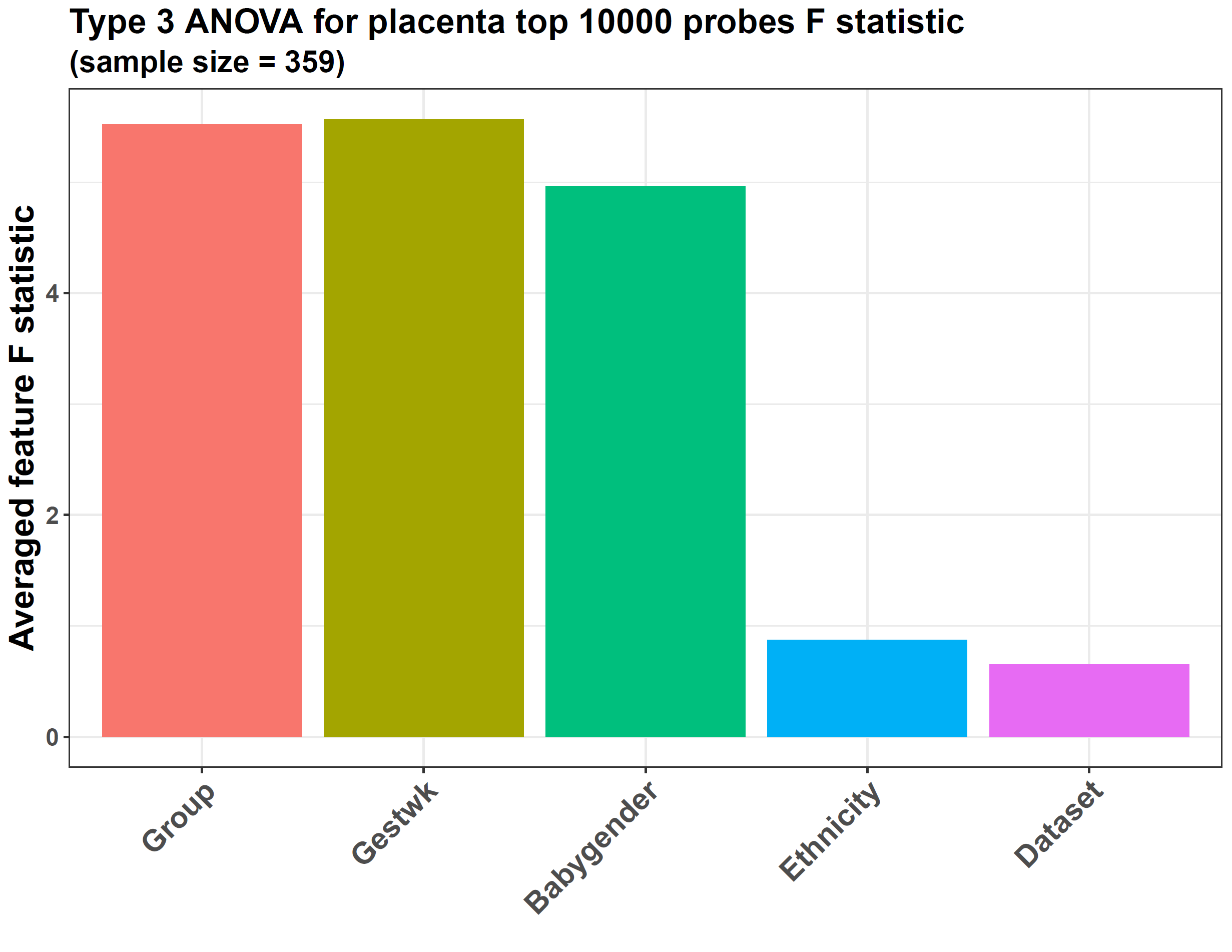

In this case, each probe in the data will go through ANOVA, and its result F statistic, MSS (mean sum of the square), and p-value will be contained in the slot varianceres in the list anovares returned by *featuresampling*. The plot generated shows the average F statistic across all the input probes.

From the plot, the preeclampsia/control sample group has the largest averaged F value, implying this dataset is suitable for checking the DNAm difference between the sample groups. However, the variances from gestational age and baby gender are also high, with an F value > 1, especially gestational age, which strongly correlated with preeclampsia because preeclampsia led to preterm delivery, so the samples always had small gestational week values *[4]*. Hence, gestational age and baby gender are confounding factors for preeclampsia analysis, and their variance needs to be adjusted. On the other hand, the variable Dataset only has a very small F statistic, meaning the batch difference from different original datasets is small, validating the batch adjustment during preprocessing of the data.

### Ensemble-based *limma* analysis

Next, we use the function *difffeatures* to call the differential DNAm probes between preeclampsia and control groups, with gestational week and baby gender adjusted.

This function performs normal *limma* or ensemble-based *limma* to achieve this purpose. The advantage of ensemble-based *limma* is that it can improve the power of *limma* for imbalanced data, which is the case here because the placenta dataset has a much larger control group than the preeclampsia one (258 samples/101 samples).

Ensemble-based *limma* uses up-down sampling to generate several base learner datasets with balanced groups, i.e., a large group will be down-sampled, and a small one will be up-sampled with SMOTE (synthetic minority over-sampling technique) so a final set with balanced groups will be generated. This process will be conducted several times to generate several such sets. Then, *limma* will be used on each of them, and their results will be ensembled to get the final. In the paper on *eOmics*, we showed a simulated experiment, and the result was that ensemble-based *limma* could generate a differential feature set much closer to the true set.

We use *difffeatures* to conduct ensemble-based *limma*. Its parameter dat accepts the betas matrix, and pddat accepts the pds data frame. The parameters responsevarname and confoundings are important because the former can define which column in pds is the target variable, and the latter can define which confoundings need to be adjusted during the regression of *limma*, and we transfer “Group” to the former, and “Gestwk” and “Babygender” to the latter. All of them correspond to the column names in pds.

We also need to set the parameter balanceadj as TRUE to trigger ensemble-based *limma*, and samplingmethod is used to define how to perform its sampling step to generate the balanced base learner sets, and we set it as “updn”. nround defines the number of base learner datasets will be included in the ensemble framework and we set it as 10.

The parameter pvalcolname is set as “adj.P.Val”, and pvalcutoff is 0.05, indicating that the differential features should have an adjusted p-value < 0.05. Because our data values here are DNAm beta values, we set the parameter isbetaval as TRUE. In this case, the log2FC result of *limma* indicates the beta value difference between the groups, and because we set absxcutoff as 0, any features with an absolute log2FC > 0 or < 0 can be considered for differential feature selection, which means we do not have a requirement on the log2FC of differential features.

For the parameter removereg, we transfer the previous DMR data frame gestdmrs to it. Also, we indicate the features in the DNAm matrix are probes via the parameter featuretype so that any probes located in the gestational age-relevant DMRs will be excluded, and the confounding effects from gestational age, the most powerful confounding factor, can be further removed.

If the parameter plot were set as TRUE, a volcano plot would be generated to show the differential features, and the characters transferred to titleprefix will be shown in the plot title. The labelnum parameter indicates how many top features in the up-regulated and down-regulated groups will be labeled in the plot. It can also be set as NULL so that no feature names will be labeled. We set it as 5 here. The parameter annotextsize can control the font size of these feature labels.

```
metadiffres <- difffeatures(dat = betas,
                            pddat = pds,
                            responsevarname = "Group",
                            confoundings = c("Gestwk", "Babygender"),

                            balanceadj = TRUE,
                            samplingmethod = "updn",
                            nround = 10,
                            seed = 2022,

                            pvalcolname = "adj.P.Val",
                            pvalcutoff = 0.05,
                            isbetaval = TRUE,
                            absxcutoff = 0,

                            threads = 4,
                            removereg = gestdmrs,
                            featuretype = "probe",

                            plot = TRUE,
                            titleprefix = "Placenta UP-DN balanced",
                            labelnum = 5,
                            titlesize = 16,
                            textsize = 16,
                            annotextsize = 6)
```

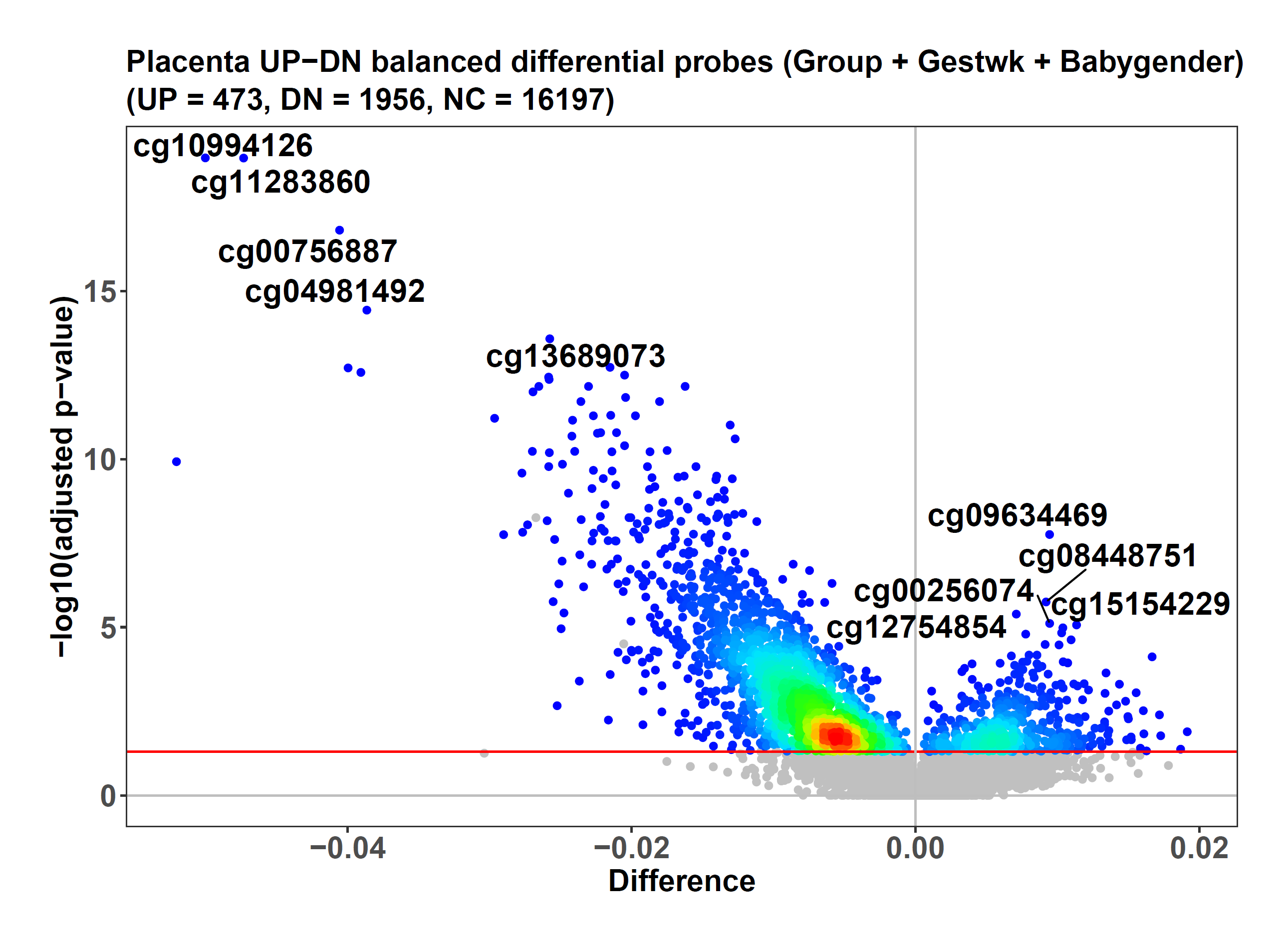

The result metadiffres is a list. Its first slot, named “limmares”, shows the ensemble-based *limma* results for each DNAm probe, including the adjusted p-value, log2FC, etc. The other slot, “plotdat”, is the organized data for generating the volcano plot.

```
head(metadiffres$limmares)
#>                  logFC      P.Value    adj.P.Val remove
#> cg10994126 -0.05001975 5.957686e-24 1.109679e-19  FALSE
#> cg11283860 -0.04732483 1.205246e-23 1.122445e-19  FALSE
#> cg00756887 -0.04056393 2.518236e-21 1.563489e-17  FALSE
#> cg04981492 -0.03864391 8.018262e-19 3.733704e-15  FALSE
#> cg13689073 -0.02575811 7.123208e-18 2.653537e-14  FALSE
#> cg06940574 -0.02151252 6.035314e-17 1.873563e-13  FALSE

head(metadiffres$plotdat)
#>               adj.P.Val        logFC   Type   color labeled
#> cg10590292 5.528198e-09 -0.026734151 NonSig #C0C0C0   FALSE
#> cg04662594 1.768817e-06 -0.010515252 NonSig #C0C0C0   FALSE
#> cg21022247 1.842171e-05 -0.009799880 NonSig #C0C0C0   FALSE
#> cg00002426 3.127396e-05 -0.020548759 NonSig #C0C0C0   FALSE
#> cg14706739 4.502128e-03 -0.003030759 NonSig #C0C0C0   FALSE
#> cg22628694 5.013027e-02  0.004135846 NonSig #C0C0C0   FALSE
```

From the volcano plot, *difffeatures* calls 473 hyper and 1956 hypomethylated DNAm probes for the preeclampsia group. The colorful dots represent the significantly differential probes, and the grey dots are the insignificant ones. However, we can see a few dots with an adjusted p-value < 0.05 but are still colored as grey, and these are the dots located in the gestational age-relevant DMRs, so even if their p-values reach the cutoff, they are still judged as insignificant ones. We check these probes.

```
subset(metadiffres$plotdat, adj.P.Val < 0.05 & Type == "NonSig")
#>               adj.P.Val        logFC   Type   color labeled
#> cg10590292 5.528198e-09 -0.026734151 NonSig #C0C0C0   FALSE
#> cg04662594 1.768817e-06 -0.010515252 NonSig #C0C0C0   FALSE
#> cg21022247 1.842171e-05 -0.009799880 NonSig #C0C0C0   FALSE
#> cg00002426 3.127396e-05 -0.020548759 NonSig #C0C0C0   FALSE
#> cg14706739 4.502128e-03 -0.003030759 NonSig #C0C0C0   FALSE
```

Hence, because the regression step of *limma* adjusted the confoundings before the DMR filtering, only 5 probes here need to be further filtered by the DMRs, so its effect is limited and unnecessary, and the default value of the removereg parameter is NULL, meaning the features do not need to go through this genomic region filter step.

If the parameter balanceadj were set as FALSE, normal *limma* would be performed by the function.

### Mediation-coupled *WGCNA* analysis

Next, we conduct *WGCNA* analysis on the data. However, before that, we use our function *probestogenes* to compress the DNAm probe beta values to genes.

betasgene <- probestogenes(betadat = betas, group450k850k = c("TSS200", "TSS1500", "1stExon"))

The parameter group450k850k means that for a specific gene, the probe located in its TSS200, TSS1500, and 1stExon regions will be selected, and their mean value will be the gene’s beta value.

Then, we use another function, *diffwgcna*, to perform the *WGCNA* analysis. Many parameters of it are the same as *difffeatures*, such as dat, pddat, responsevarname, etc. Because *diffwgcna* uses *limma* to find the significantly differential *WGCNA* modules and the differential features within each module, the parameters responsevarname and confoundings are needed for the regression step of *limma*. In addition, removereg and featuretype can also be used, so the genes within confounding-relevant genomic regions will be excluded initially.

We previously also tried combining *WGCNA* analysis with an ensemble framework. However, the result showed that it could not improve the performance, so this function only performs the normal *WGCNA* method to call the modules. Although it also has the parameter balanceadj, it is used to control whether the *limma* and mediation analysis should be performed in an ensemble manner, not the *WGCNA* module calling.

We set the parameter topvaricancetype as “sd” and topvaricance as 5000, meaning we will perform this *WGCNA* analysis on the top 5000 most variable genes in the data (the ones with the highest standard deviation).

We transfer the vector seq(1, 20, 1) to powers, so grid search will be used to select the optimal soft-thresholding power from 1 to 20 for *WGCNA*, and the desired minimum scale-free topology fitting index is 0.8 because we set another parameter rsqcutline as 0.8.

The parameter mediation is important because it controls whether mediation analysis needs to be performed during the *WGCNA* pipeline. For each *WGCNA* module, it constructs mediation models to test 2 causal directions. One is “module->module gene->preeclampsia”, and the other is “preeclampsia->module gene->module”. Hence, we can determine whether the disease causes the module changes or the module drives the disease. The parameters responsevarname and confoundings will also be used when constructing the mediation models.

Other parameters such as plot, titleprefix, pvalcolname, etc., are the same as *difffeatures*. The only difference is the additional parameter diffcutoff. It is similar to absxcutoff, but it defines the cutoff on module eigengene difference to select differential modules, and absxcutoff defines the feature log2FC cutoff to select differential features within the modules.

This *WGCNA* analysis is time-consuming, and the running below can be skipped.

```
wgcnares <- diffwgcna(dat = betasgene,
                      pddat = pds,
                      responsevarname = "Group",
                      confoundings = c("Gestwk", "Babygender"),
                      removereg = gestdmrs,
                      featuretype = "gene",

                      topvaricancetype = "sd",
                      topvaricance = 5000,

                      balanceadj = TRUE,
                      samplingmethod = "updn",
                      nround = 10,
                      seed = 2022,

                      powers = seq(1, 20, 1),
                      rsqcutline = 0.8,

                      mediation = TRUE,

                      plot = TRUE,
                      titleprefix = "Placenta",
                      titlesize = 17,
                      textsize = 16,
                      annotextsize = 5,

                      pvalcolname = "adj.P.Val",
                      pvalcutoff = 0.05,
                      isbetaval = TRUE,
                      absxcutoff = 0,
                      diffcutoff = 0,

                      threads = 4)
```

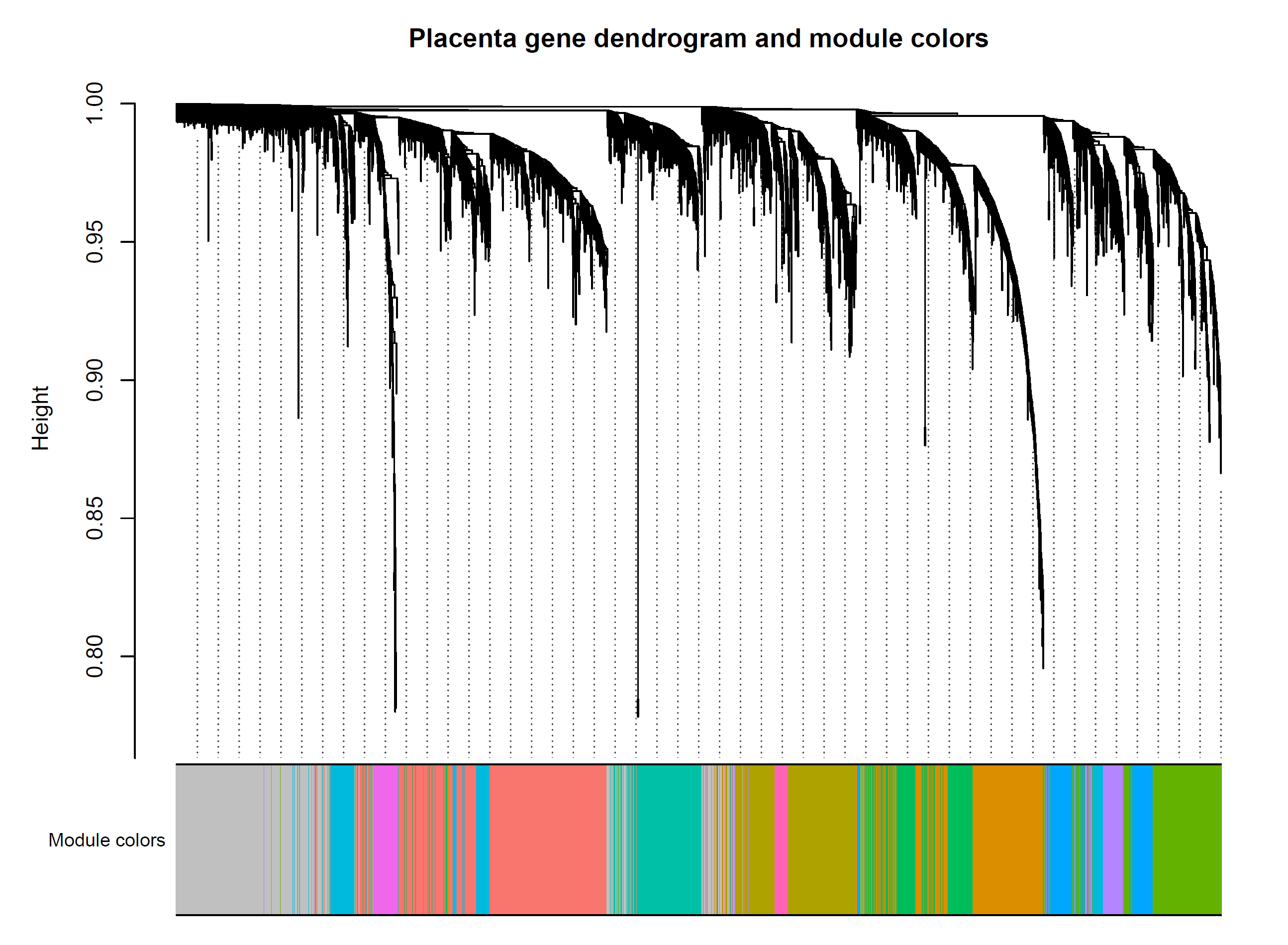

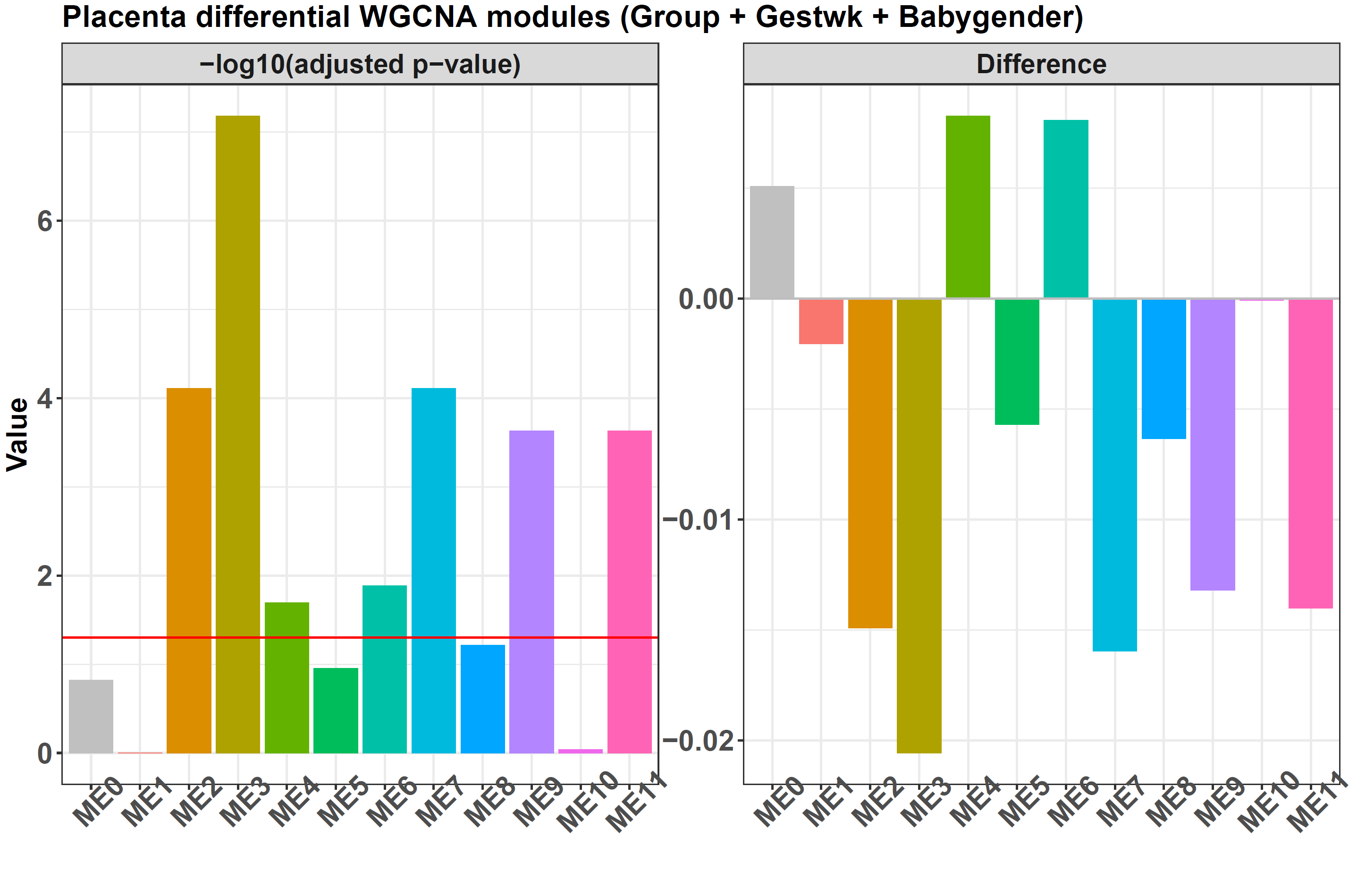

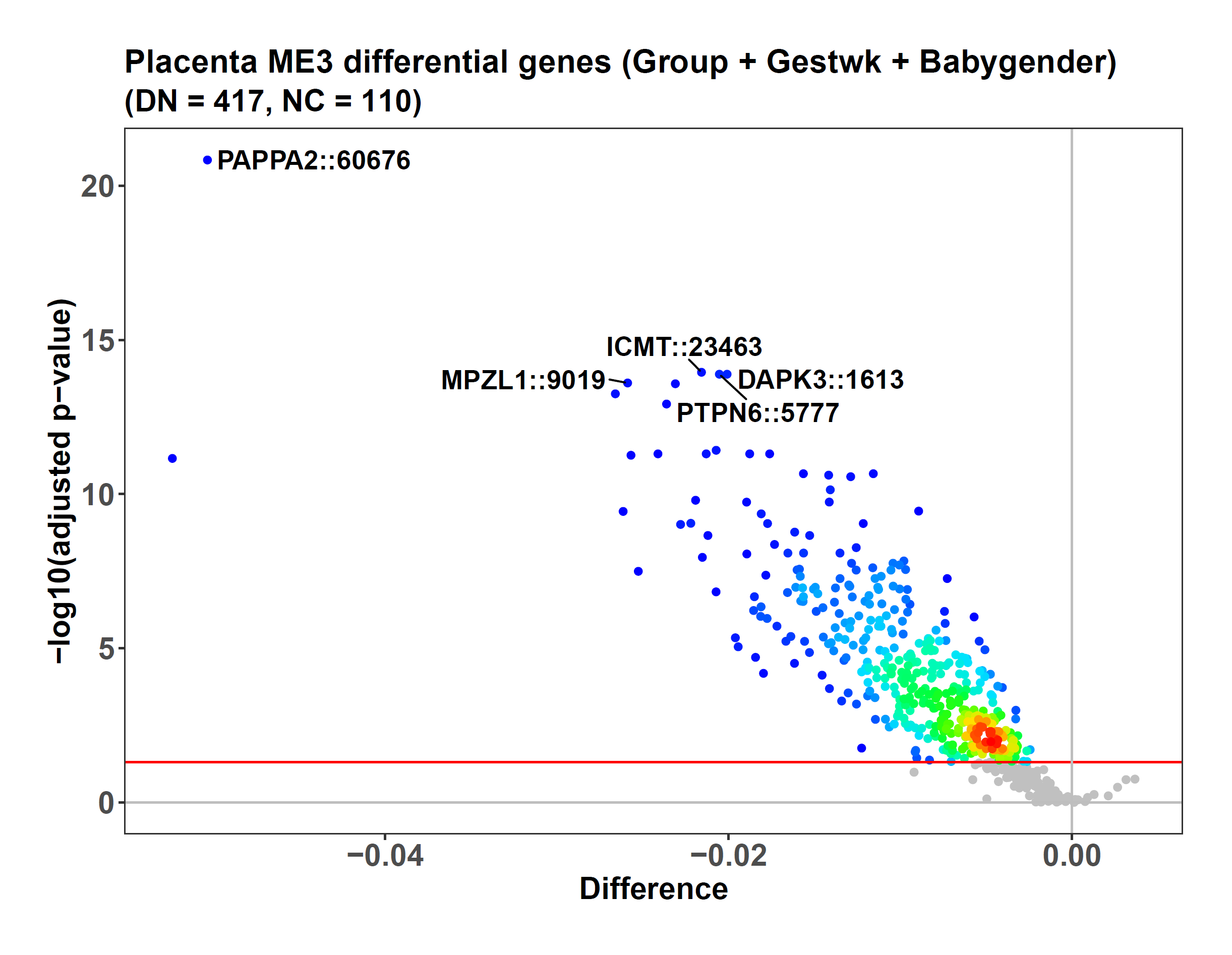

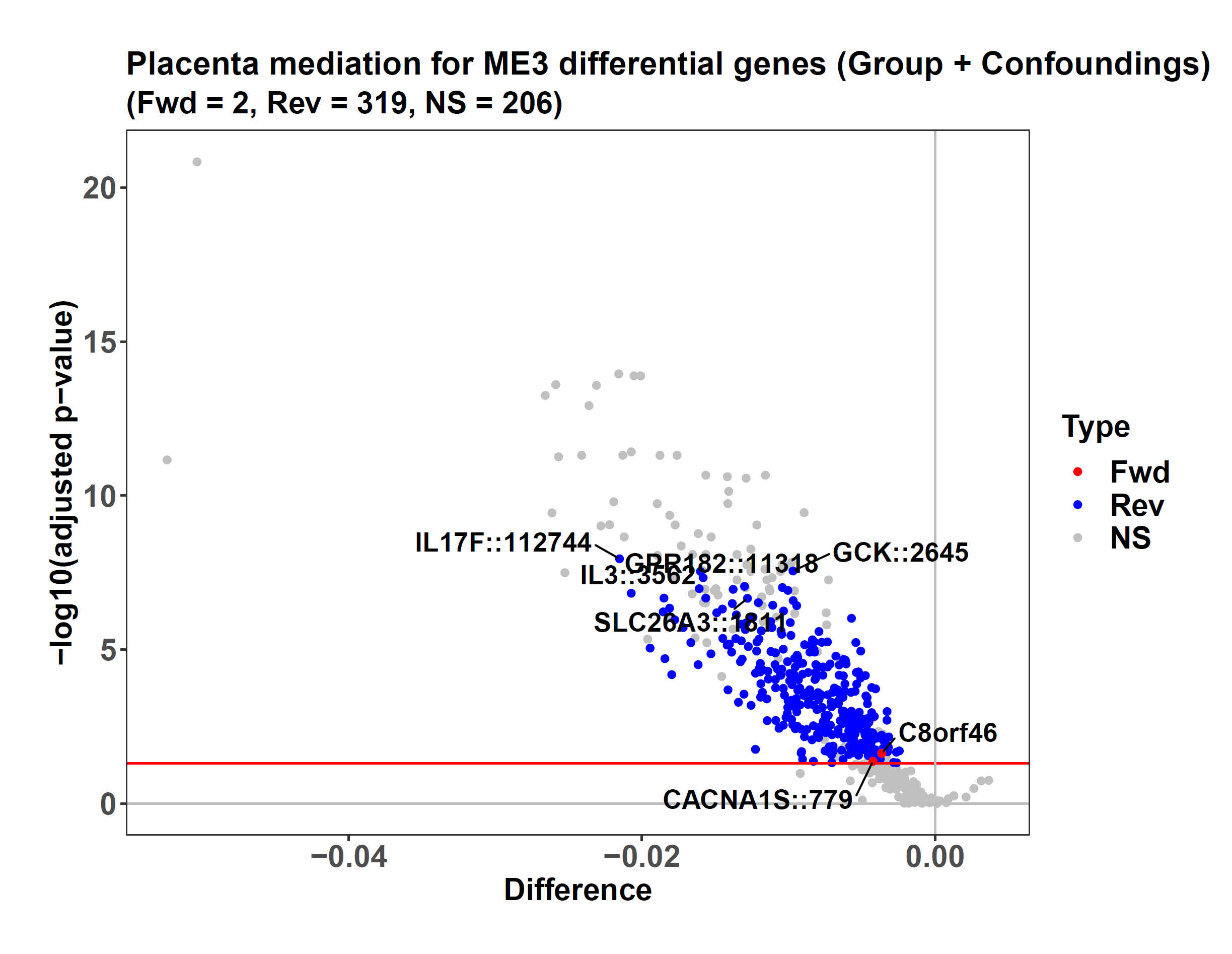

The result wgcnares is a list containing several slots. The one named “limmares” is the ensemble-based *limma* result on all the *WGCNA* modules, and the bar plot generated by the function is based on this result. It shows the data contain 11 modules from ME1 to ME11, and 7 have significantly differential eigengenes between preeclampsia/control groups.

In addition, ensemble-based *limma* also selects the differential features within each module. For example, the volcano plot of ME3 shows 417 of its 527 genes are hypo-methylated in preeclampsia compared to the control. The slot “melimmares” contains the *limma* results within each significantly differential module.

Because the parameter mediation is set as TRUE, the mediation results for the differential modules are also returned, and the slot “mediationres” records them. Correspondingly, each module gets an additional volcano plot to show its mediation result. For example, in the second volcano plot for ME3, the 319 blue dots are genes mediating the causal direction of “ME3->ME3 gene->preeclampsia” (the reverse direction), and the red dots mediate the direction of “preeclampsia->ME3 gene->ME3” (the forward direction). The y-axis and x-axis show the -log10(adjusted p-value) and preeclampsia/control beta value difference when screening the differential genes with *limma*.

The 319 genes show an association with inflammatory stress in preeclampsia, such as IL3, IL17F, and GPR182, which are the top genes mediating the effects of preeclampsia on ME3. Among them, IL3 and IL17F are cytokines for inflammatory responses *[5,6]*, and GPR182 is a G protein-coupled receptor interacting with chemokines CXCL10, 12, and 13 *[7]*. Because these genes’ causal direction is from preeclampsia to ME3, the conclusion is that inflammation is not the cause, but the result, of this disease.

Meanwhile, CACNA1S and C8orf46 (VXN) are the only 2 genes mediating the causal direction from ME3 to preeclampsia. CACNA1S has expression in the placenta *[8,9]*, and encodes the alpha-1S subunit of the dihydropyridine receptor, a calcium channel responsible for vascular contraction. Given that preeclampsia is a pregnancy hypertension disease, the causal effect of CACNA1S on it is clear. Moreover, this gene is the target of amlodipine besylate, a small-molecule drug for preeclampsia treatment, further supporting that the inference here is reasonable.

At the same time, the other gene, C8orf46, is largely unknown. However, it is still reported to work together with CDKN1B (P27KIP1) *[10]*, a critical gene for mediating trophoblast invasion *[11]*. On the other hand, it is generally agreed that insufficient invasion of placental trophoblasts is a driver of preeclampsia *[12]*. Hence, C8orf46 may cause this disease by influencing trophoblast invasion.

### PCA/CCA-based clustering and ensemble-based elastic net classification

Finally, we use our package to perform some machine-learning tasks. We first use the function *multiCCA* to cluster the 101 preeclampsia samples to see if any subtypes exist. The function *multiCCA* uses CCA to convert multi-omics data into a compressed one and then performs k-means to get the clustering result. On the single-omic here, the CCA becomes PCA for the DNAm data of the 101 samples.

```
#Extract the DNAm probe data for the 101 preeclampsia samples
prepds <- subset(pds, Group == "Preeclampsia")
row.names(prepds) <- 1:nrow(prepds)

prebetas <- betas[, prepds$sampleid, drop = FALSE]

#Clustering

presubtyperes <- multiCCA(dats = list(prebetas),

                          k = 2,
                          consensus = 1,

                          seednum = 2022,
                          threads = 4,
                          plot = TRUE,
                          titlefix = "Preeclampsia",
                          titlesize = 18,
                          textsize = 16)
```

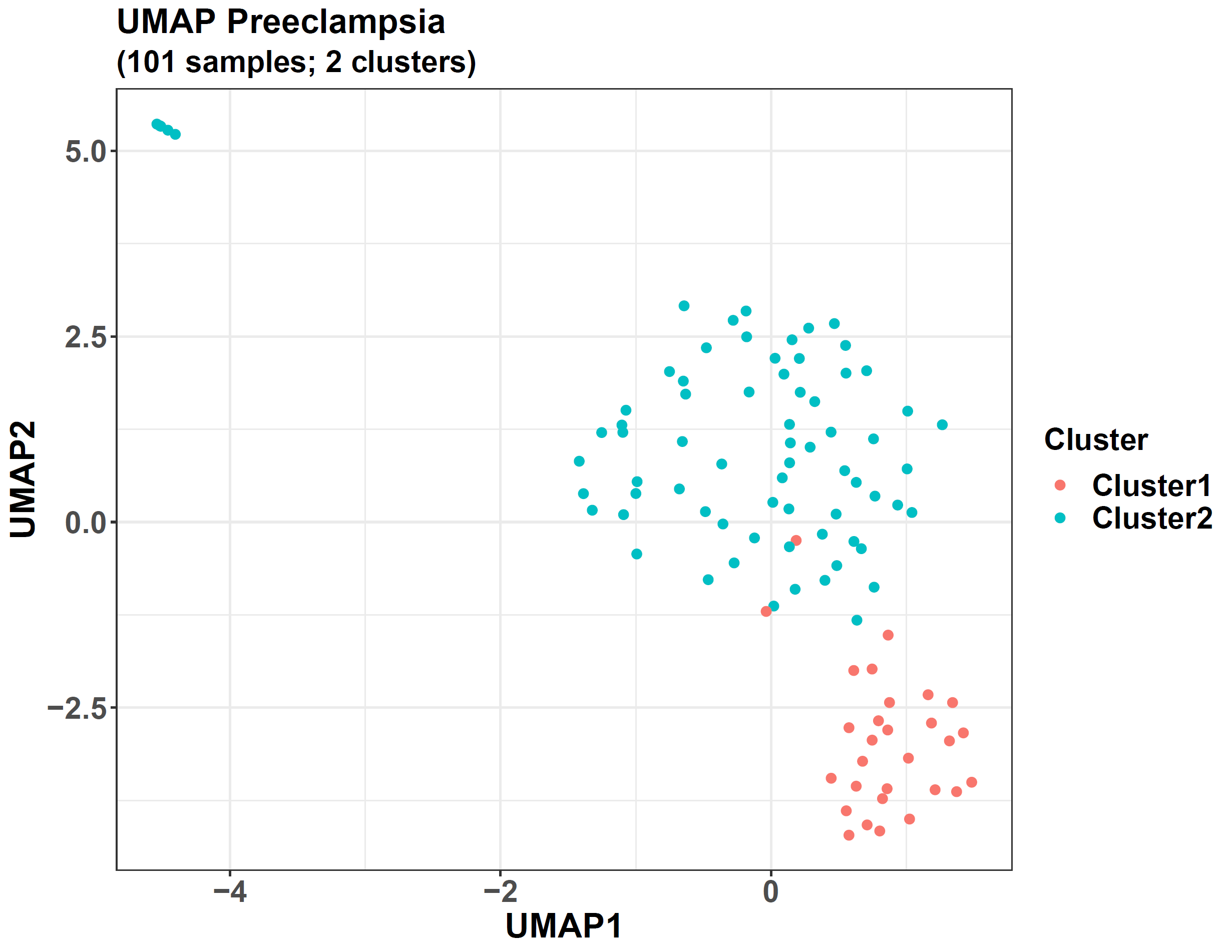

The parameter dats accepts a list with each element as the data of one omic, and because our current task is single-omic, we transfer a one-element list with the preeclampsia DNAm data.

The parameter k defines the cluster number for the k-means step, and if the optimal cluster number is not known, a vector, such as c(2, 3, 4) can be transferred, so for each number in it, the k-means step will use it as the cluster number, and the optimal one can be defined by the internal validation indices calculated by *multiCCA*. From our previous experiment, the optimal cluster number for this preeclampsia dataset is 2, so we can directly use 2 here and do not need to transfer more candidate numbers. Another parameter, consensus, is used for consensus clustering, and if we set it as 100, then for each cluster number, consensus clustering will be conducted, and its feature shuffling will be performed 100 times. However, we set it as 1 here, and a normal clustering will be conducted without the consensus process.

Because the parameter plot is TRUE, the clustering results will be plotted.

The result presubtyperes is a list, and its slot “ivis” records 3 internal validation indices for each candidate cluster number.

```
presubtyperes$ivis
#> $`k = 2`
#>        silhouette calinski_harabasz    davies_bouldin
#>         0.2622399        36.6521374         1.3709824
```

We only transfer one number to the parameter k, so the Silhouette, Calinski, and Davies indices are only calculated for the cluster number of 2. If more candidate numbers are transferred, these indices will also be calculated for their clustering results, and the optimal cluster number should have higher Silhouette and Calinski indices but a smaller Davies index, which means that its intra-cluster samples are closer to each other, and the inter-cluster sample distance are larger.

The slot “kreses” contains the cluster labels for samples, and its sub-slot “k = 2” is for the cluster number of 2.

```
head(presubtyperes$kreses$`k = 2`)
#> GSM1892055 GSM2674426 GSM2674432 GSM2589560 GSM2674433 GSM1892048
#>          2          1          2          2          2          1

table(presubtyperes$kreses$`k = 2`)
#>
#>  1  2
#> 29 72
```

Hence, 29 samples belong to Cluster1 (subtype1), and 72 belong to Cluster2 (subtype2). The function also returns their UMAP plot, and the 2 clusters are separated well, supporting that 2 subtypes exist in the preeclampsia data.

After getting the subtype labels, we can train a classifier to classify the samples using the function *omicsclassifier*. It combines elastic net with the balanced ensemble framework to improve the performance on imbalanced data, such as the subtype1/subtype2 case here.

```
#Make the sample labels from the clustering result
subtypes <- paste0("Subtype", presubtyperes$kreses$`k = 2`)

#Classification
presubtypeclassifierres <- omicsclassifier(dats = list(prebetas),
                                           truelabels = subtypes,

                                           balanceadj = TRUE,
                                           nround = 10,

                                           alphas = c(0.5),
                                           nfold = 5,

                                           seednum = 2022,
                                           threads = 4,

                                           plot = TRUE,
                                           prefixes = c("Preeclampsia (UP-DN)"))
```

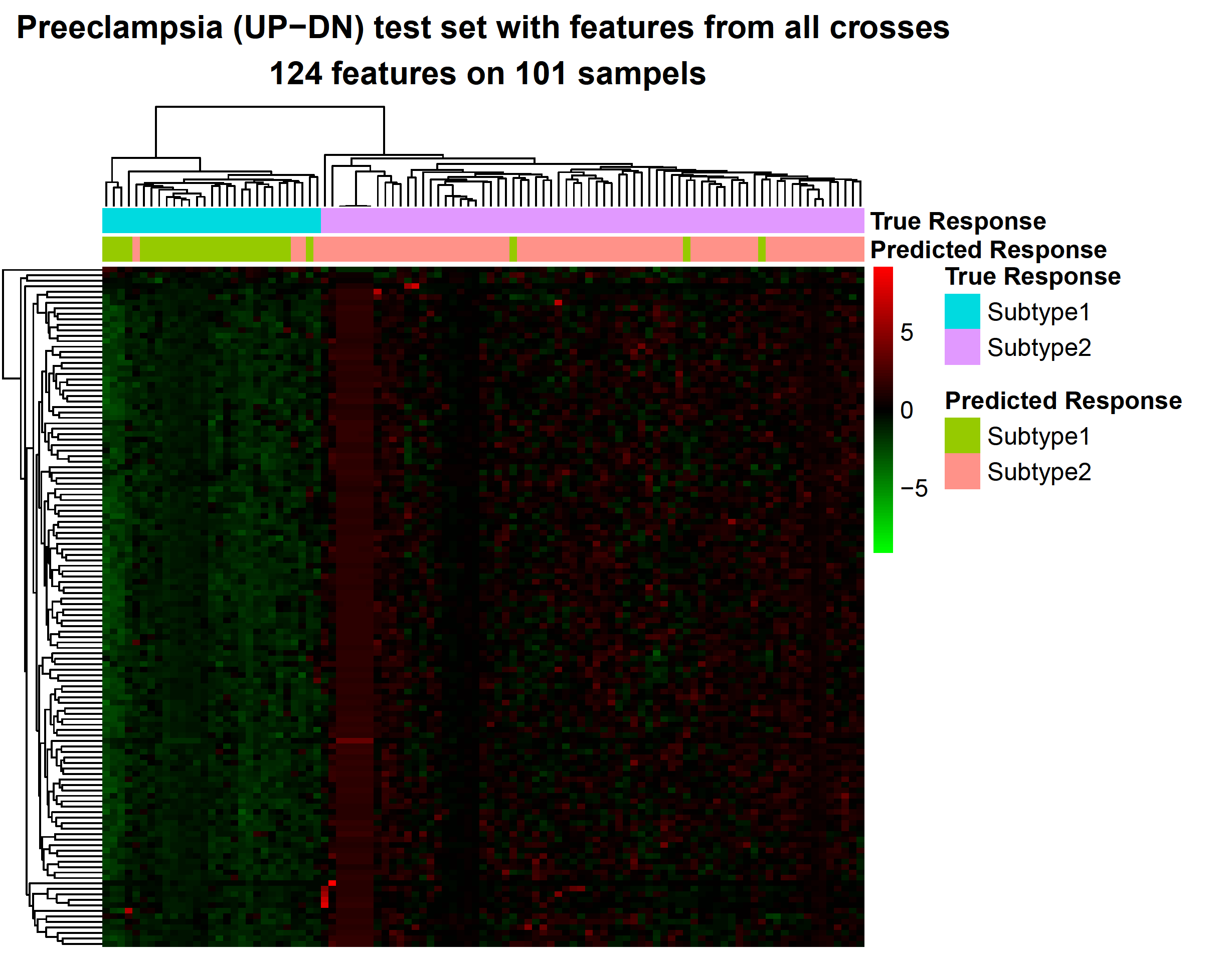

omicsclassifier can handle both single-omic and multi-omics data, and in this case, we transfer the DNAm data of the preeclampsia samples to its parameter dats, similar to that in multiCCA. The parameter truelabels needs the subtype labels from the clustering result.

Then, balanceadj triggers an ensemble-based elastic net model, the same as difffeatures and diffwgcna.

alphas accepts a number between 0 and 1, the alpha parameter of the elastic net model, controlling the balance of L1 and L2 penalties. Another parameter, nfold, indicates whether a cross-fold validation should be performed, and we set it as 5, so a 5-fold cross-validation will be used to evaluate the performance of the ensemble-based elastic net model. If it were set as NULL, cross-fold validation would not be performed, and only a model trained on the whole dataset would be returned.

The parameter plot is set as TRUE, so the function will generate some plots to show the final results.

The result presubtypeclassifierres contains a slot named “cvtestcomps”, showing the predicted and true labels for the testing data during the 5-fold cross-validation.

```
head(presubtypeclassifierres$cvtestcomps)
#>            Prediction     True
#> GSM1892031   Subtype2 Subtype2
#> GSM1892044   Subtype2 Subtype2
#> GSM1892047   Subtype1 Subtype1
#> GSM1892048   Subtype1 Subtype1
#> GSM1892055   Subtype2 Subtype2
#> GSM1892056   Subtype2 Subtype2

#Number of correct classifications
sum(as.character(presubtypeclassifierres$cvtestcomps$Prediction) == as.character(presubtypeclassifierres$cvtestcomps$True))
#> [1] 94

#Accuracy
sum(as.character(presubtypeclassifierres$cvtestcomps$Prediction) == as.character(presubtypeclassifierres$cvtestcomps$True))/nrow(presubtypeclassifierres$cvtestcomps)
#> [1] 0.9306931
```

From this slot, we can see that 94 of the 101 samples are classified correctly, so the 5-fold cross-validation accuracy is 0.931. It is also shown by the heatmap returned by the function, where each column represents one sample, and each row represents one DNAm probe selected by the model. The entries are beta values after scaling across the samples.

On the other hand, if the parameter balanceadj were set as FALSE, a normal elastic net model would be trained, and our previous results showed its accuracy was 0.881, weaker than the ensemble-based elastic net model. It is because the balanced base learners of that model help improve the model performance.

The result presubtypeclassifierres also contains a slot named “mod”, which is the final model trained from the whole dataset, and it can be transferred to another function *pairedensemblepredict*, which will use the model to predict other external samples.

In addition to the functions above, *eOmics* also has other functions to perform structure-based gene functional enrichment (*diffwgcna* and *topoenrich*), multi-omics regulatory analysis (*lassomediation*), etc., and more details can be found in our package document and the paper of this package.

### References

1. Ritchie ME, Phipson B, Wu D et al. limma powers differential expression analyses for RNA-sequencing and microarray studies, Nucleic Acids Res 2015;43:e47.
2. Zhang B, Horvath S. A General Framework for Weighted Gene Co-Expression Network Analysis, Statistical Applications in Genetics and Molecular Biology 2005;4.
3. Kuleshov MV, Jones MR, Rouillard AD et al. Enrichr: a comprehensive gene set enrichment analysis web server 2016 update, Nucleic Acids Research 2016;44:W90-W97.
4. Davies EL, Bell JS, Bhattacharya S. Preeclampsia and preterm delivery: A population-based case–control study, Hypertension in Pregnancy 2016;35:510-519.
5. Dougan M, Dranoff G, Dougan SK. GM-CSF, IL-3, and IL-5 Family of Cytokines: Regulators of Inflammation, Immunity 2019;50:796-811.
6. Chang SH, Dong C. IL-17F: Regulation, signaling and function in inflammation, Cytokine 2009;46:7-11.
7. Le Mercier A, Bonnavion R, Yu W et al. GPR182 is an endothelium-specific atypical chemokine receptor that maintains hematopoietic stem cell homeostasis, Proceedings of the National Academy of Sciences 2021;118:e2021596118.
8. Bernucci L, Henriquez M, Diaz P et al. Diverse Calcium Channel Types are Present in the Human Placental Syncytiotrophoblast Basal Membrane, Placenta 2006;27:1082-1095.
9. Zhao Y, Pasanen M, Rysa J. Placental ion channels: potential target of chemical exposure, Biology of Reproduction 2022.
10. Moore KB, Logan MA, Aldiri I et al. C8orf46 homolog encodes a novel protein Vexin that is required for neurogenesis in Xenopus laevis, Developmental Biology 2018;437:27-40.
11. Nadeem L, Brkic J, Chen YF et al. Cytoplasmic mislocalization of p27 and CDK2 mediates the anti-migratory and anti-proliferative effects of Nodal in human trophoblast cells, Journal of Cell Science 2013;126:445-453.
12. Kaufmann P, Black S, Huppertz B. Endovascular Trophoblast Invasion: Implications for the Pathogenesis of Intrauterine Growth Retardation and Preeclampsia, Biology of Reproduction 2003;69:1-7.
